# Visualizing the mechanism of quinol oxidation and inhibition of a *bd*-type oxidase using cryo-EM

**DOI:** 10.1101/2025.10.10.681541

**Authors:** Tijn T. van der Velden, Kanwal Kayastha, Famke Pelser, Steffen Brünle, Lars J. C. Jeuken

## Abstract

Cytochrome *bd* is a prokaryotic terminal oxidase recognized as an antibiotic target against various pathogens. Despite its critical role in respiration, failure to capture the mechanism of enzyme catalysis and inhibition prohibits structure guided drug discovery. Here, we present cryo-electron microscopy structures of *Escherichia coli* cytochrome *bd*-I in monomeric and dimeric forms, along all stages of quinol turn-over and in an inhibitor-bound state. We identify a dynamic Q-loop lid that undergoes a disorder-to-order transition upon substrate binding to the dimer, completing the active site and enabling catalysis. Structure-guided mutagenesis confirms Tyr243 and Arg298 as essential catalytic residues unique to long Q-loop oxidases, highlighting evolutionary divergence from short Q-loop variants. Inhibition by Aurachin D triggers refolding of the active site, occluding substrate access via a conserved Asp239-mediated mechanism. The structural and mechanistic insights presented here establish a comprehensive framework, opening new ways for drug discovery.

## Introduction

The bacterial respiratory chain has emerged as a promising target for next-generation antibiotics due to its essential role in ATP generation(*1*). Disruption of this pathway has proven effective in eliminating pathogens resistant to conventional therapeutics, as demonstrated by the clinical success of bedaquiline, an ATP synthase inhibitor approved for the treatment of multidrug-resistant tuberculosis(*2*). Among the terminal oxidases, cytochrome *bd* (cyt *bd*) is characterized by an exceptionally high oxygen affinity and ability to confer resistance to oxidative stress and tolerance against diverse antibiotics (*3*–*8*). This enables pathogens to sustain respiration under hostile conditions encountered during infection. Notably, cyt *bd* is absent in eukaryotes, making it an attractive target for selective antimicrobial development against pathogens such as *Mycobacterium tuberculosis*(*9*), *Salmonella enterica*(*10*), pathogenic *Escherichia coli*(*11*), and *Staphylococcus aureus*(*12*).

Cyt *bd* catalyzes the reduction of molecular oxygen to water using electrons from the quinone pool. During this turnover, cyt *bd* releases the chemical protons associated with quinol oxidation to the periplasm while capturing cytoplasmic protons for oxygen reduction, thereby contributing four protons per oxygen reduced to the proton motive force required for ATP synthesis (Fig. 1A,B). The canonical enzyme architecture comprises two core subunits, CydA and CydB, and in some species, auxiliary subunits such as CydS, CydX, and CydH(*13, 14*). CydA harbors the three heme cofactors, *b*_*558*_, *b*_*595*_, and *d*, as well as the putative quinone binding site near the Q-loop, a flexible periplasmic region adjacent to heme *b*_*558*_(*13*). Cyt *bd* oxidases are classified into long and short Q-loop subfamilies based on the presence of a C-terminal Q-loop extension, though the functional implications of this divergence remain unresolved(*15, 16*).

**Figure 1.**
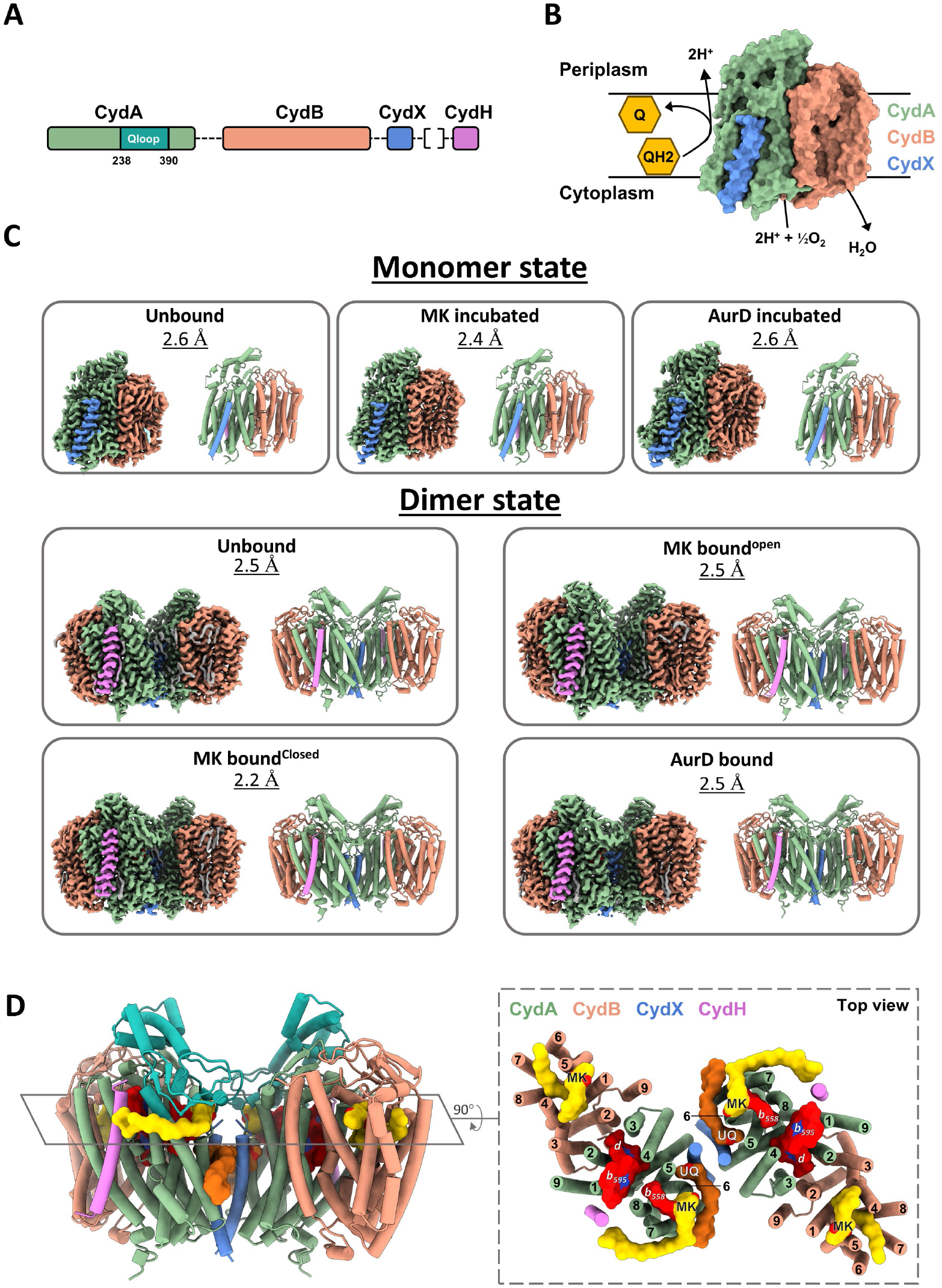
Overview of the structures and function of *Ecbd*. (**A**) Genomic organization of the *Ecbd* subunits. Subunit CydH originates from a different operon than the other *Ecbd* subunits. (**B**) Schematic oxidase activity of *Ecbd*. (**C**) Overview of the different *Ecbd* states resolved using cryo-EM. Both colored volume maps and cylinder models are shown with their respective resolution. (**D**) Structural overview of the *Ecbd* dimer in the MK bound state with the Q-loop highlighted. CydA is shown in green, with the Q-loop in cyan, CydB in orange, CydX in blue, CydH in purple, MK in yellow and UQ In brown. The right shows a cross section of the *Ecbd* dimer with a total of 40 transmembrane helices, highlighting all identified MK (yellow) and UQ (dark orange), as well as heme groups (red).

*E. coli* cyt *bd-I* (*Ecbd*), representative of the long Q-loop subfamily, consist of the four subunits CydA, CydB, CydX and CydH. The available substrate pool in *E. coli* consists of three quinone subtypes, ubiquinone (UQ), demethylmenaquinone (DMK) and menaquinone (MK). The composition of this quinone pool adapts based on the environmental oxygen concentration, switching from predominantly UQ to MK under micro-aerobic conditions when *Ecbd* is expressed(*17*). Despite extensive characterization(*13, 18, 19*), structural and mechanistic insights into the turnover and inhibition mechanism of cyt *bd* have remained elusive.

To gain insights into quinone turnover and inhibition of *bd*-type oxidases, we have resolved Cryo-EM structures of *Ecbd* in its unbound, MK bound and inhibitor bound state. We identified both monomeric and dimeric forms of *Ecbd*, with the latter exhibiting increased catalytic activity. The dimeric form revealed three discrete quinone binding sites per protomer. In the active site, located near heme *b*_*558*_, MK was observed to occupy a cavity formed by a disorder-to-order transition of the Q-loop lid. Unexpectedly, the active site undergoes a structural rearrangement upon binding of the inhibitor Aurachin D, occluding the quinone from binding to its pocket. Utilizing these structures, we highlight the full catalytic cycle of *Ecbd*, and underline critical structural features required for quinol turnover and inhibition.

## Results

### Purification and structure determination of *Ecbd*

*Ecbd* was expressed in MB43 cells and purified using affinity chromatography followed by gel filtration using the detergent LMNG. To gain insights into the turnover mechanism of *Ecbd*, we determined cryo-EM structures of the (A) unbound, (B) menaquinone bound in the open and (C) closed, as well as (D) inhibitor bound states (Table 1, SI Fig. 1-4). All structures were resolved in LMNG, except for the menaquinone-bound sample, which was reconstituted into phosphatidylcholine (PC) nanodiscs doped with 5% menaquinone-9 to enhance local substrate concentration.

Our cryo-EM analysis identified previously uncharacterized dimeric forms of *Ecbd*, similar to the homologue *E. coli cyt bd-II*(*20*), for (A), (B), (C), and (D) at 2.2-2.5 Å overall resolution. Additionally, on the same grids, the canonical monomeric architecture for sample (A), (B), and (C) were resolved at 2.4-2.6 Å overall resolution. However, all three monomeric structures show an identical state, with no additional ligand binding being observed other than a structural quinone in CydB.

Consistent with previous reports(*13*) *Ecbd* adopts a heterotetrameric architecture that consists of the subunits CydA, CydB, CydX and CydH (Fig. 1A). CydA forms the catalytic core, harboring the three heme cofactors (*b*_*558*_, *b*_*595*_, and *d*) and the putative quinone-binding Q-loop required for oxidase activity (Fig. 1B). In contrast to previous reports, our high resolution cryo-EM map indicates that the heme *d* hydroxychlorin γ-spirolactone exists in the trans rather than the cis orientation (SI Fig. 5). This trans orientation is further confirmed by prior NMR and IR analysis of the isolated heme(*21*), and positions the spirolactone group opposite the bound dioxygen molecule. Close examination of prior Cyt *bd* density maps indicates the possibility of a more general trans heme d orientation within the *bd*-oxidase family, although unambiguous assignment of the chirality remains challenging due to limitations in resolution.

The *Ecbd* dimer consists of 20 transmembrane helices existing in a symmetrical architecture around the CydX dimer interface (Fig. 1D). CydX forms hydrophobic contacts with CydA helix 1, 5, and 6 to connect the two protomers together. To confirm the physiological relevance of both monomer and dimeric states, *Ecbd* was extracted using styrene–maleic acid lipid particles (SMALPs). Cryo-EM imaging followed by 2D classification showed clear populations consisting of dimeric and monomeric particles, indicating the existence of both oligomeric states *in vivo* (SI Fig. 6). Both oligomeric states contained tightly bound phospholipids at the protein periphery and dimer interface (SI Fig. 7). Lipids lacking discernible headgroups were modeled as phosphatidic acid (PA), suggesting either headgroup flexibility or phospholipid heterogeneity.

### Quinone Binding and Electron Transfer in the *Ecbd* Dimer

The dimer interface of *Ecbd* is formed by CydX in conjunction with coordinated phosphatidylglycerol (PG) lipids (Fig. 2A,B). These PG lipids also interact with the CydA Q-loop hydrogen bonding between Arg298 and the lipid carbonyl group. The homodimeric CydX– CydX interface is stabilized by a methionine–aromatic motif (Met1^CydX^–Trp2^CydX^) and a leucine zipper along the helical axis (Fig. 2B).

**Figure 2.**
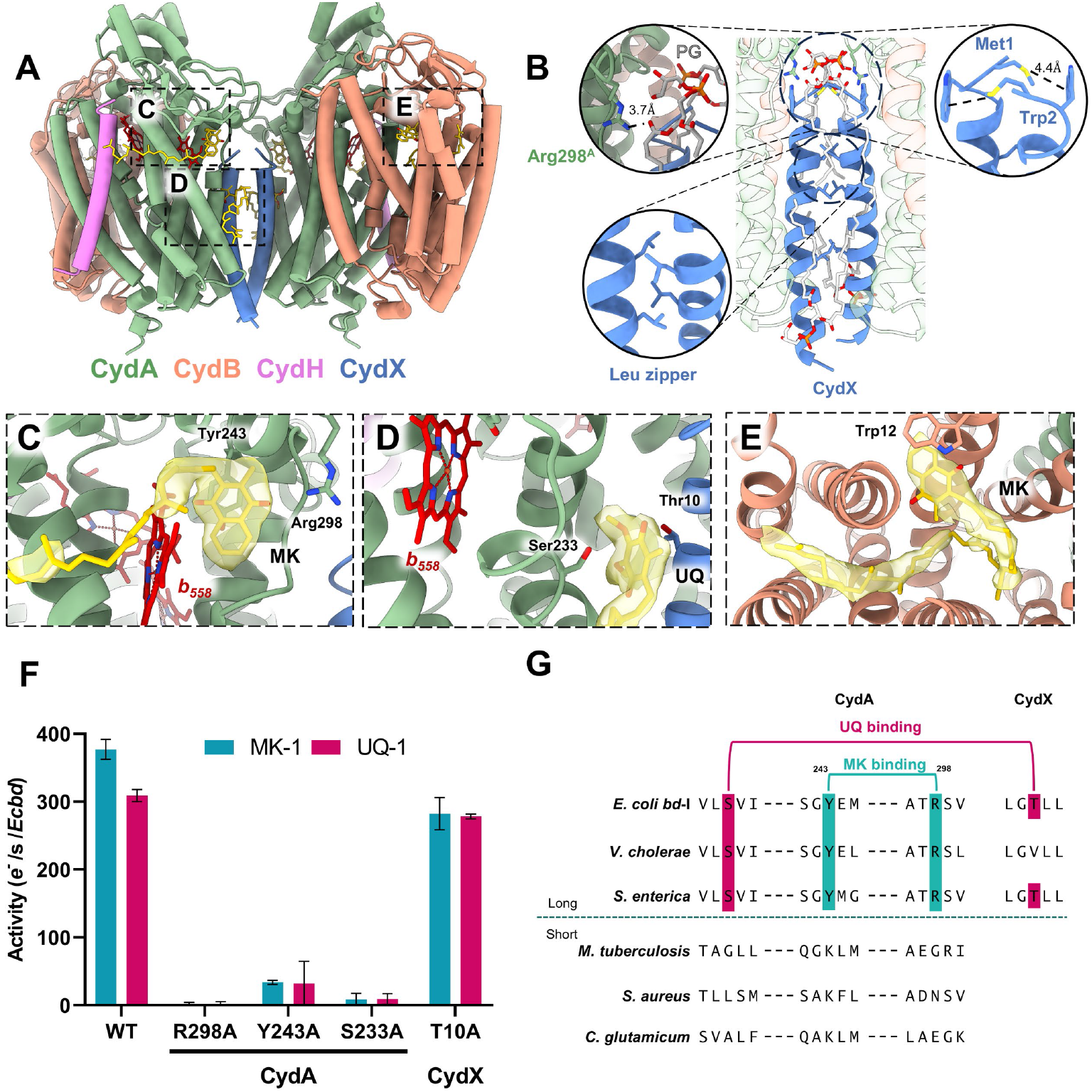
The *Ecbd* dimer contains three distinct quinone binding sites per protomer. (**A**) Overview of the *Ecbd* dimer and the quinone binding pockets. (**B**) The *Ecbd* dimer interface is formed by subunit CydX via Met-Trp interactions, and a leucine zipper. Additionally, an PG phospholipid forms a salt bridge with CydA Arg298 further stabilizing the interface. (**C**) The Q binding site near heme *b*_*558*_, showing a bound MK stabilized by Tyr243 and Arg298. (**D**) The UQ binding site wedged between subunit CydA and CydX. The UQ is stabilized by interactions with CydA Ser233 and CydX Thr10. (**E**) The quinone binding pocket in CydB with a bound MK, stabilized by π–π stacking with CydB Trp12. The quinone headgroup is situated outside of electron transfer distance from the heme chain. (**F**) oxidase activity of *Ecbd* WT and quinone binding site mutants (n=3) (**G**) Representative sequence alignment of long- and short – Q-loop Cyt*bd* variants indicating conservation of Y243^CydA^ and R298^CydA^ in the long Q-loop family (extended sequence alignment in SI Fig. 11).

Within the dimer, each *Ecbd* protomer contains three distinct quinone-binding pockets. The primary quinone binding site, located beneath the Q-loop, accommodates a well-resolved menaquinone (MK) headgroup (Fig. 2C). This shallow, hydrophobic cavity is positioned 6.7 Å from the heme *b*_*558*_ iron center, enabling fast electron transfer. The MK isoprenoid tail is only partially resolved, indicating its flexibility during quinone binding and turnover. The MK headgroup is stabilized by π–π stacking with Tyr243^CydA^ (4.5 Å) and hydrogen bonding with Arg298^CydA^ (3.4 Å), both of which facilitate semiquinone stabilization and proton transfer during catalysis. This site is strongly implicated as the quinol oxidation center, as mutagenesis of either Tyr243^CydA^ or Arg298^CydA^ to alanine (Y243A^CydA^, R298A^CydA^) abolished enzymatic activity with both MK and ubiquinone (UQ), confirming its role as the principal active site (Fig. 2F). The arrangement of the three hemes in CydA follows a triangular topology, as previously described (13), with the MK-bound structure facilitating mapping of the full electron transfer route from quinone binding site to the oxygen reduction center (Fig. 3A). Although no proton release pathway could be identified unambiguously, the protons released during quinol oxidation could transfer to the periplasm via the closely positioned Asp239^CydA^ and Glu240^CydA^ pair at the membrane interface, or transfer via a series of structured water molecules to the conserved Lys252^CydA^ (SI Fig. 8). The associated electrons are transferred sequentially from this MK site via heme *b*_*558*_ to *b*_*595*_ (14.9 Å) and finally to heme *d* (11.2 Å), where oxygen reduction occurs in two cycles of quinol oxidation (Fig. 3A).

**Figure 3.**
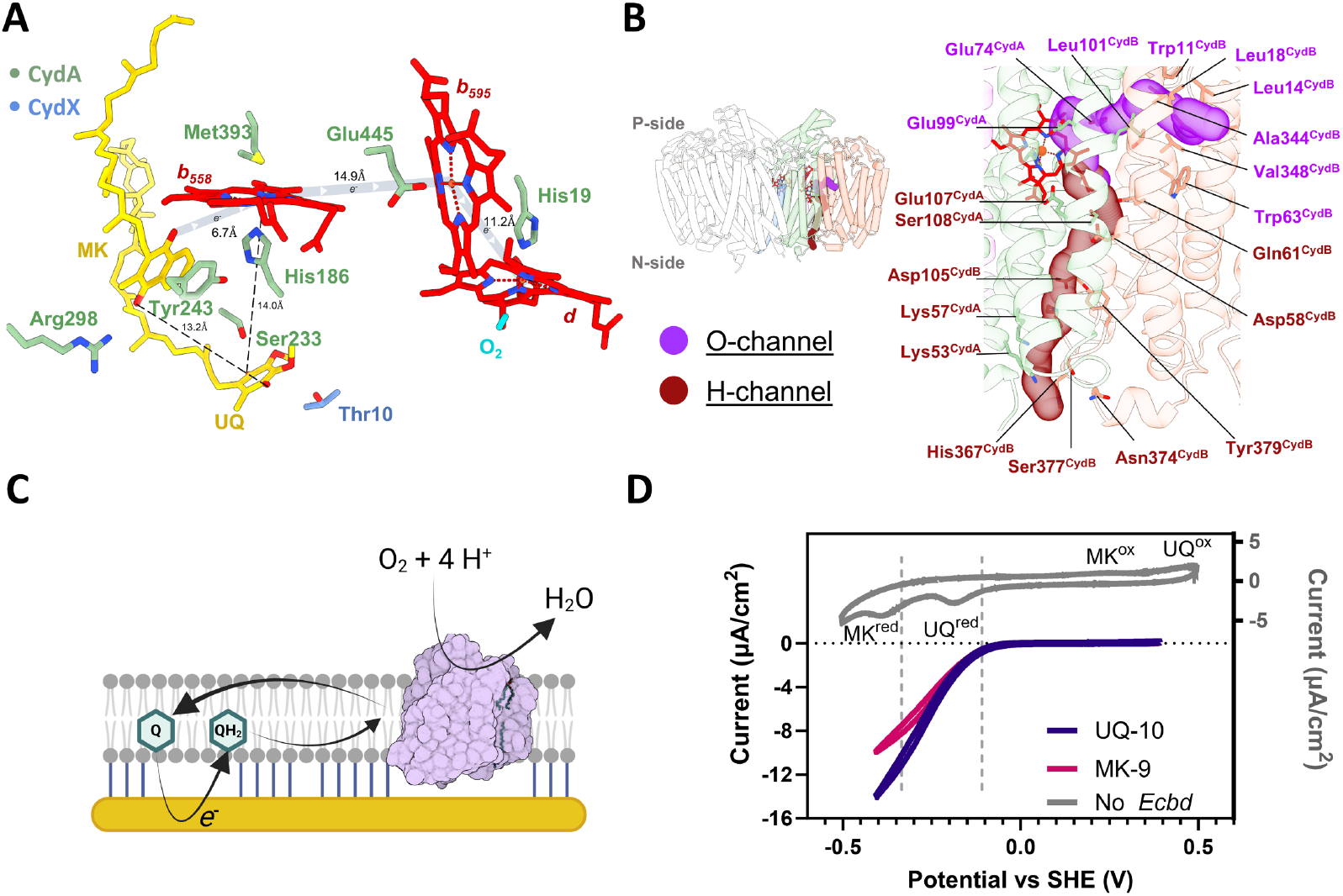
The electron transfer, proton, and oxygen pathways. (**A**) Proposed electron transfer routes from the two separate quinone binding sites (**B**) Residues lining of the O and H-channel leading towards the heme *d* oxygen reduction center. (**C**) Overview of the tBLM electrochemical system, where the quinone pool is directly reduced at the gold electrode to initiate *Ecbd* turnover. (**D)** Cyclic voltammograms of the tBLM system. Inset in grey shows a model membrane containing both MK (1 mol%) and UQ (1 mol%) without *Ecbd*. The reduction and oxidation peaks of UQ and MK are indicated. The SAM negative control (black), *Ecbd* tBLMs containing only UQ-10 purple), or MK-9 (pink) showing the same onset potential at the reduction of UQ.

A second quinone binding site is located at the CydA–CydX interface and contains a co-purified ubiquinone molecule (Fig. 2D). This UQ is coordinated via a hydrogen-bonding network involving Ser233^CydA^ and Thr10^CydX^ of the same protomer. Positioned 14 Å from the heme *b*_*558*_ iron center and 13.2 Å from the primary MK site, this second site lies within electron transfer distance, with potential for both direct electron transfer and indirect electron transfer into the heme chain. However, mutation of S233^CydA^ to alanine in the second quinone binding pocket (S233A) in CydA also fully inactivates *Ecbd*, while mutation T10^CydX^ to alanine (T10A) remains active (Fig. 2F). Although the activity of T10A^CydX^ strongly suggests the second quinone binding site is not an active site, we further investigate the abolished activity of the S233A^CydA^ mutant. Notably, the S233A^CydA^ mutant exhibited no heme reduction after incubation with dithionite, in contrast to the WT and T10A^CydX^ mutant (SI Fig. 9), implicating S233^CydA^ has a structural role rather than a role in oxidation of the ubiquinol substrate. Thus, while this site is capable of binding quinones tightly, retaining UQ even after incubation with excess MK, it likely does not function as a site for quinol oxidation.

A third quinone site, typically occupied by UQ in CydB(*13, 19*), was found to contain MK in the MK-incubated structure (Fig. 2E). Although the density for this MK is less defined, indicating either heterogeneity or flexible binding, the headgroup appears repositioned near Trp12^CydB^, differing from the unbound structure. Despite this rearrangement, the MK headgroup remains 30 Å from the heme *d* center and lacks nearby residues to stabilize a semiquinone, indicating a structural rather than catalytic role. Functional assays confirmed that quinone exchange at this site does not impact enzymatic activity, precluding any allosteric effects (SI Fig. 10).

Sequence alignment across *bd*-type oxidases revealed conservation of the Tyr243^CydA^ or Arg298^CydA^ residues exclusively within the long Q-loop subfamily (Fig. 2G, SI Fig. 11), indicating an evolutionary divergence in active site architecture between short and long Q-loop variants.

### Functional role of the quinone pool in *Ecbd* regulation

While *Ecbd* is active with both MK and UQ(*22*), we aimed to assess if the content of the quinone pool provides any regulation on *Ecbd* activity. To assess this putative allosteric regulation by the available quinone pool, we employed a tethered bilayer lipid membrane (tBLM) electrochemical system. This system enables direct electron transfer from a gold electrode to membrane-embedded quinones (Fig. 3C)(*23*), and has previously been used to study a variety of respiratory enzymes(*24*–*26*). To form the tBLM, *Ecbd* was reconstituted in liposomes doped with either MK-9 or UQ-10 as native-like substrates.

Formation of the tBLM was confirmed by a characteristic drop in surface capacitance(*27*). Reduction of the quinone pool and subsequent quinol oxidation by *Ecbd* was monitored via cyclic voltammetry (CV), which can distinguish UQ and MK based on their redox potentials. CV measurements revealed catalytic currents for both the UQ-10 and MK-9 *Ecbd* samples with identical onset potentials and similar catalytic currents, indicating that the co-purified UQ in the MK sample is redox-active and capable of supporting *Ecbd* turnover (Fig. 3D). Whether this activity originates directly from the CydA–CydX site or reflects quinone exchange at the primary active site remains to be determined. We conclude that *Ecbd* is promiscuous regarding the quinol pool and not allosterically regulate by either UQ or MK.

### Proton and oxygen delivery to the oxygen reduction center

Following the reduction of heme *d*, molecular oxygen and protons need to be guided towards the oxygen reduction site to complete the catalytic cycle. Using Mole 2.5(*28*) we identified two distinct channels converging at the heme *d* site, previously deemed the proton and oxygen channels (Fig. 3B)(*13*). The oxygen channel (O-channel) originates between helices 1 and 9 of CydB and runs parallel to the membrane. The channel is lined with hydrophobic residues to facilitate the preferential entry of molecular oxygen. Near heme *d*, the channel is terminated by two glutamate residues providing hydrogen bonding during oxygen reduction.

The H-channel originates at the CydA–CydB interface and extends perpendicularly from the cytoplasmic side toward heme *d*, and is lined with titratable residues forming a putative proton wire. Our predictions using PROPKA of residue pKa suggest proton transfer involving Lys53^CydA^ (pKa 9.01), Lys57^CydA^ (pKa 7.45), Asp105^CydB^ (pKa 4.53), Asp58^CydB^ (pKa 8.14), Glu107^CydA^ (pKa 9.47), and Glu74^CydA^ (pKa 11.07). The high pKa of Lys53 allows for fast proton extraction from the cytoplasm to increase turnover rates. Among the channel residues, Asp105 emerges as the most acidic and has been previously implicated as critical in proton transfer(*29*). Notably, Glu107, positioned 7.8 Å from the heme *d* iron center, exhibits a pKa consistent with the proposed terminal proton donor identified by Janczak et al.(*29*) and is highly conserved across CydA homologues (>94% sequence conservation). These findings support a model in which Lys53 initially extracts protons from the cytoplasm, after which Glu107 serves as the primary proton donor for oxygen reduction, completing the proton delivery pathway.

### Closing of the Q-loop lid completes the active site and enhances catalytic efficiency

Despite exposure to high concentrations of menaquinone (MK), no density is observed near the Q-loop in monomeric *Ecbd*, indicating persistent conformational flexibility. This precludes reliable modeling of residues Glu239–Gly306^CydA^ in the *Ecbd* monomer. Structural comparison between monomeric and dimeric *Ecbd* revealed a rotation of Trp2^CydX^ upon dimerization, facilitating hydrogen bonding with CydA Asp239^CydA^ (Fig4. A,B). This interaction anchors the Q-loop segment spanning Asp239–Thr247^CydA^, positioning Tyr243^CydA^ optimally for quinol capture. Together, this suggests that the dimeric state is more conducive to MK binding and, hence, MK oxidation.

*Ecbd* monomers and dimers were separated using gel filtration to probe the activity of the individual states. Each state showed to be stable in detergent solution by SEC-MALS, enabling kinetic evaluation (SI Fig. 11). Catalytic activity was assessed using a coupled assay in which quinones were reduced by an excess of NADH dehydrogenase(*22*). The oxygen consumption rate was determined using an oxygraph and recalculated into *Ecbd* activity at 4 electrons per oxygen reduced. The observed structural stabilization in the *Ecbd* dimer correlates with a twofold increase in catalytic turnover rate relative to the monomer, without a significant change in substrate *K*_*M*_ (Fig. 4C). These findings suggest that in the monomer, transient Q-loop remodeling is required to form a functional active site, rendering this conformational transition rate-limiting. In contrast, the dimeric enzyme maintains a partially-formed, catalytically competent active site.

**Figure 4.**
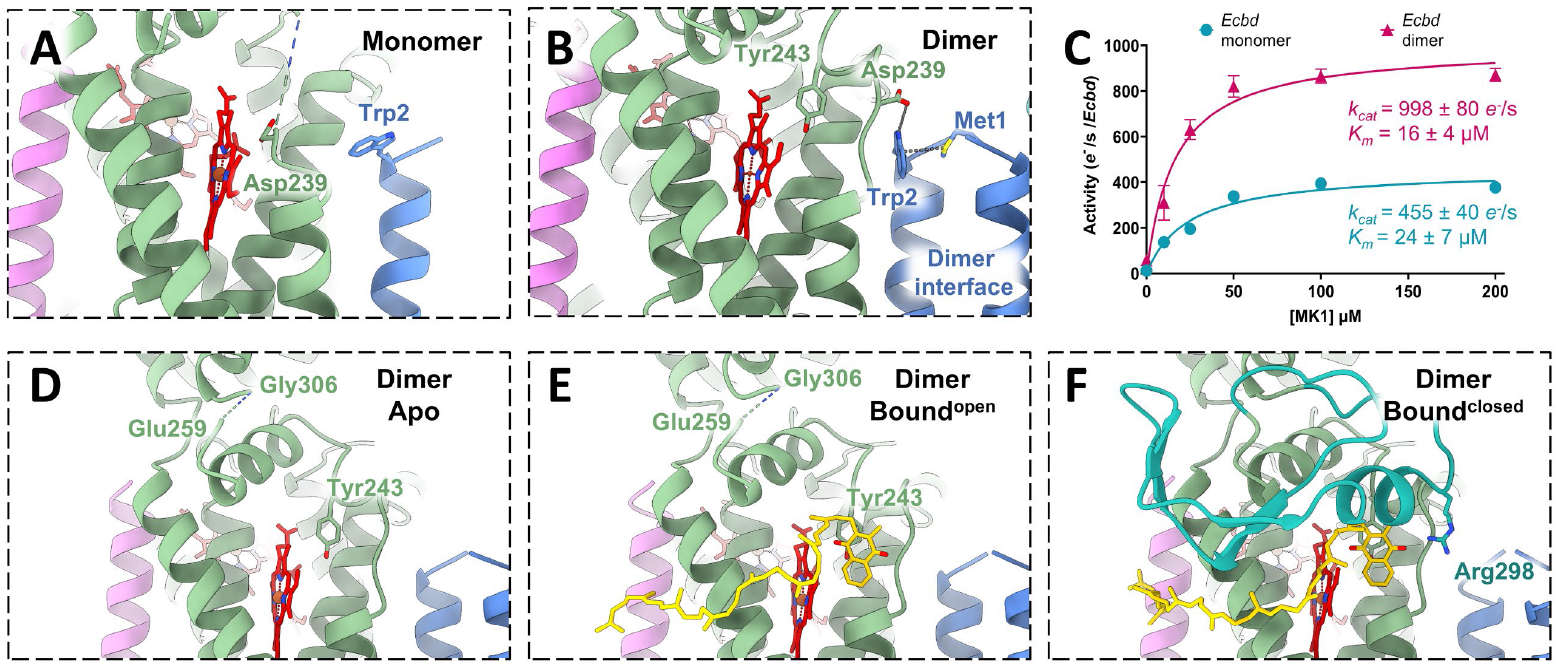
Q-loop stabilization increases the turnover rate in the *Ecbd* dimer. (**A**) Selected view of the quinone binding pocket in the *Ecbd* monomer (**B**) Menaquinone binding at the Q-loop active site. The quinone is coordinated by Tyr243 and Arg298. (**C**) Michaelis Menten kinetics of the *Ecbd* monomer and *Ecbd* dimer (n=3). (**D**) unbound *Ecbd* dimer. (**E**) *Ecbd* dimer in the MK bound^open^ state. (**F**) *Ecbd* dimer in the bound^closed^ state. CydA is represented in green, with the Q-loop lid in dark green. CydH is represented in violet, with CydX in blue.

Although the Asp239–Thr247^CydA^ segment is stabilized in the dimer, the remainder of the Q-loop remains disordered in the unbound state (Fig. 4D). This precludes modelling of the amino acid stretch between Glu259^CydA^ and Gly306^CydA^, hereafter referred to as the Q-loop lid. This flexible region remains unstructured even after initial MK engagement by Tyr243, showing (Fig. 4E). Only in this MK bound state can the Q-loop lid undergo a structural rearrangement to stabilize the fold, enabling confident modelling of side chains and revealing the complete active site architecture. The Q-loop lid wraps around the quinone binding site before looping back onto the main body of CydA, mirroring the fold found in *Ecbd-II* and *M. tuberculosis* cyt *bd*(*20, 30*). The rearrangement of the Q-loop lid allows for hydrogen bonding between Arg298^CydA^ and the quinone headgroup, completing the active site for quinol turnover (Fig. 4F).

### Inhibition by Aurachin D triggers active site refolding

Aurachin D (AurD), a menaquinone analog derived from *Stigmatella aurantiaca*, is a potent and specific inhibitor of cyt *bd* oxidases(*31*). To elucidate its mechanism of action, we resolved cryo-EM structures of both monomeric and dimeric *Ecbd* in the presence of AurD. While no density for AurD was observed in the monomer, the dimeric structure revealed clear binding near the Q-loop site which allows for unambiguous modeling of the AurD binding pose (Fig. 5A).

**Figure 5.**
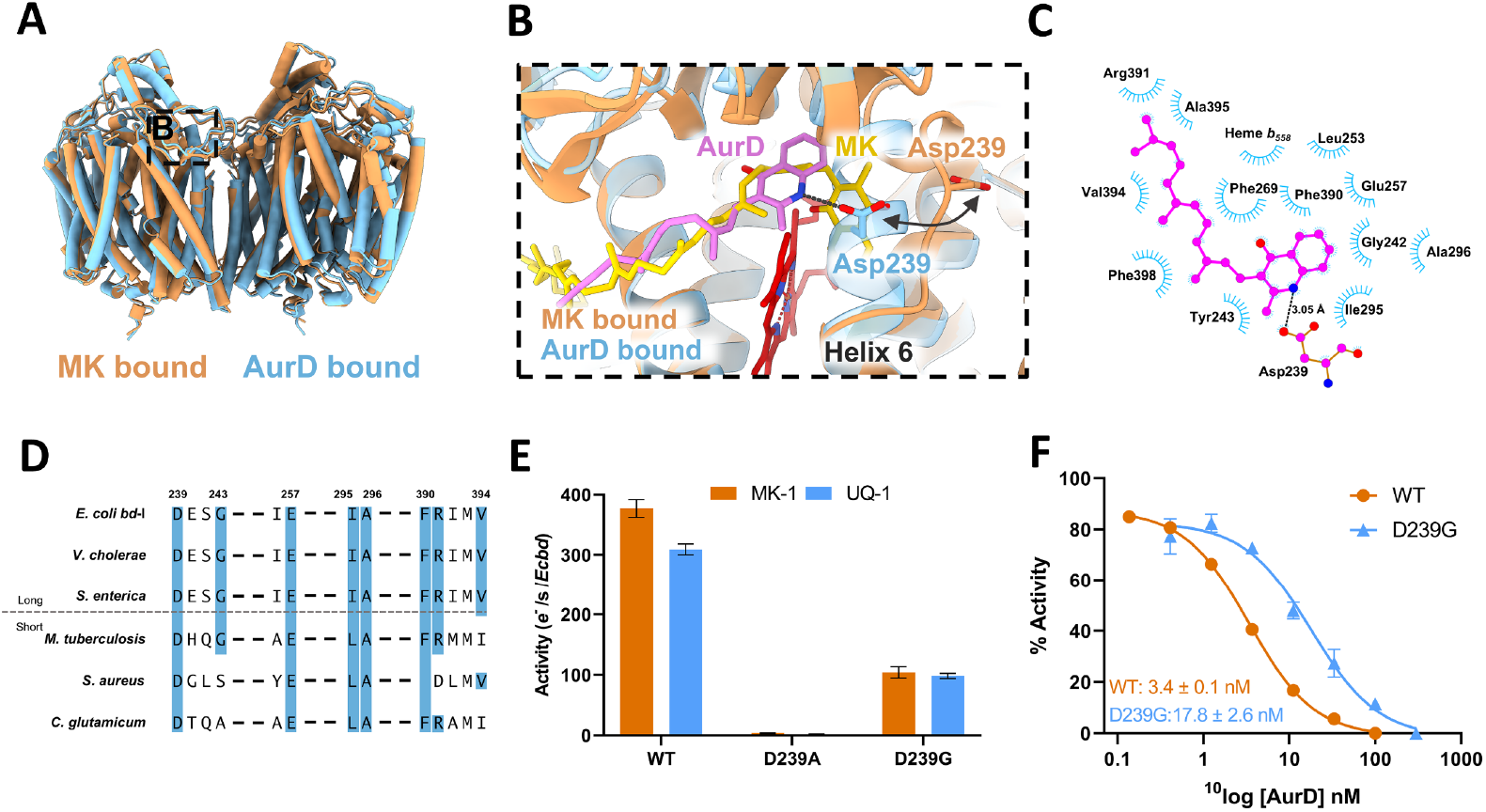
Inhibitory mechanism of AurD. (**A**) Overview of the *Ecbd* dimer in the MK and AurD bound state. (**B**) Refolding of the *Ecbd* active site by AurD by the attraction of Asp239^CydA^. (**C**) Two dimensional representations of the interactions in the AurD binding pocket. (**D**) Conservation of the AurD binding pocket residues. (**E**) Activity of the *Ecbd* WT and D239A^CydA^ and D239G^CydA^ mutants (n=3). (**F**) IC50 of AurD against the *Ecbd* WT and D239G^CydA^ mutant (n=3).

Surprisingly, AurD does not bind in the same pocket as MK, but induces a remarkable conformational change in helix 6, converting it from a loop into an α-helix. This conformational change closes the quinone binding pocket, effectively blocking substrate access. Close examination of the refolding process indicates that Asp239^CydA^ forms a hydrogen bond to the AurD amine group and hereby pulls the loop inwards to stabilize the helical conformation (Fig. 5B). The AurD headgroup is further stabilized by hydrophobic interactions with Gly242^CydA^, Tyr243^CydA^, Leu253^CydA^, Glu257^CydA^, Phe269^CydA^, Ile295^CydA^, Ala296^CydA^, Phe298^CydA^, Phe390^CydA^, Arg391^CydA^, Val394^CydA^, Ala395^CydA^, and heme *b*_*558*_ (Fig. 5C). These interactions in the AurD binding pocket are highly conserved in all *bd* oxidases (Fig. 5D, SI Fig. 13). However, AurD binding occurs deeper into the binding pocket compared to the binding in *Ecbd*-II(*20*) (SI Fig. 14).

To confirm Asp239^CydA^ as the main effector residue for inhibition with AurD we performed mutagenesis (D239A^CydA^). Although Asp239^CydA^ lies outside the active site in the MK-bound state, its mutation to alanine abolishes enzymatic activity, resulting in the inability to support bacterial growth (Fig. 5E, SI Fig. 15), consistent with prior reports(*32*). We postulate that inactivity of the *Ecbd* D239A^CydA^ mutant results from the impairment to form the hydrogen bond with CydX Trp2 (Fig 3B), effectively closing the MK binding pocket as seen in the AurD bound state. In contrast, substitution with glycine partly preserves activity while significantly reducing AurD sensitivity (Fig. 5E,F), confirming Asp239^CydA^ as a key determinant of inhibitor binding and efficacy, as seen in *Ecbd*-II (*20*).

## Discussion

The structural and mechanistic insights presented here establish a comprehensive framework for understanding quinone turnover, electron transfer, and inhibition in *bd-*type oxidases. Prior structural studies of cyt*bd* oxidases have predominantly captured the enzyme in a monomeric state, with no resolved structures of quinone-bound complexes. A notable exception is the dimeric structure of *E. coli* cyt bd-II bound to AurD, resolved in amphipols at 3.0 Å resolution(*20*). However, the physiological relevance of this dimeric state remained uncertain, as it was speculated to be an artifact of amphipol reconstitution. Using a cryo-EM approach we were able to resolve the dimeric state of *Ecbd* in detergent and nanodiscs, and show it as a native state *in vivo* by SMALP isolation as well as the presence of native *E. coli* lipids at the dimer interface. The CydX dimer interface is well conserved, indicating common dimerization of Cyt *bd* in the species containing the CydX subunit(*33*). Interestingly, CydX is solely found within a sub-population of the long Q-loop family, showing further divergence within cyt *bd* oxidases. Similar, but not homologous, single-helical subunits are found in short Q-loop bd oxidases, such as CydS in *G. thermodenitrificans* Cyt *bd*(*14*). CydS, however, does not seem to facilitate dimer formation in AlphaFold prediction (SI Fig. 14). Additionally, CydS has a profoundly different sequence than CydX, indicating a role in stabilization rather than dimerization.

Functionally, the dimeric form of *Ecbd* exhibits significantly enhanced catalytic activity compared to the monomer, suggesting that dimerization is physiologically important for its function. In the monomeric state, the Q-loop remains disordered despite substrate availability, suggesting that active site formation is transient and rate-limiting. This is further supported by previous studies showing that deletion of the dimer-forming subunit CydX compromises *Ecbd* expression, stability, and activity(*34, 35*). We propose that dimerization serves as a regulatory mechanism, enabling *Ecbd* to become catalytically active under microaerobic conditions when it is upregulated to replace the more energetically efficient cytochrome *bo*_3_. This dynamic switch may allow *E. coli* to balance respiratory efficiency with adaptability to stress.

The *Ecbd* dimer shows quinone binding in a shallow peripheral pocket that stabilizes only the headgroup, consistent with rapid substrate exchange from the quinone pool. This primary active site, located beneath the Q-loop lid near heme *b*_*558*_, is stabilized by CydA residues Tyr243^CydA^ and Arg298^CydA^, both uniquely conserved in long Q-loop bd oxidases. This conservation underscores a mechanistic divergence from the short Q-loop subfamily. Moreover, Tyr243^CydA^ and Arg298^CydA^ are distinctly different from the previously indicated active site residues, Lys252^CydA^, Glu257^CydA^, and Glu280^CydA^ (*36*). These residues stabilize the Q-loop in the menaquinone bound form by forming hydrogen bonding with the main body of CydA, which could explain the drop in activity upon mutation(*36*).

The Q-loop undergoes a disorder-to-order transition upon quinone binding, completing the active site and enabling catalysis. This large structural rearrangement during the catalytic cycle represents substantial energetic investments to form the catalytically active state(*37*). Such large structural rearrangement during catalysis has previously been indicated to prevent irreversible binding of the substrate while presenting a binding pocket highly optimized for the transition state(*38*). This means that the closed Q-loop state represents an intermediate specifically stabilized by the semi-quinone. The binding energy associated with this state will release upon quinone formation, expelling the quinone from the active site to improve turnover rates. This mechanistic adaptation might allow Cyt *bd* to maintain high catalytic activity and low *Km*, necessary to uphold membrane potential despite displacing less protons than cytochrome bo3 or other pumping terminal oxidases(*37*).

In addition to the primary site, we identified a secondary ubiquinone-binding pocket at the CydA–CydX interface. This site is specific for UQ and mutagenesis suggests it contributes to structural stabilization of the dimer rather than direct catalysis.

A particularly striking finding is the dual role of CydA Asp239^CydA^. This residue is essential for both active site stabilization and inhibition by AurD. Despite the structural similarity between MK and AurD, AurD binding induces a remarkable refolding of helix 6 into an α-helix, preventing substrate access. Interestingly, this α-helical conformation mirrors that observed in all solved structures of short Q-loop bd oxidases (SI Fig. 16)(*30, 39, 40*), where the AurD binding pocket, including Asp239^CydA^, is likewise conserved. Despite the sequence and structural conservation of the binding pocket in *M. tuberculosis* cyt *bd*, no densities have been observed for AurD in the presence of vast molar excesses(*30*). To probe a similar binding poses of AurD in short Q-loop *bd* oxidases, we performed docking in the *M. tuberculosis* cyt *bd* Q-loop. Despite the conservation of the pocket, no similar binding pose was observed (SI Fig. 17). This raises the possibility that short Q-loop *bd* oxidases undergo structural rearrangement of the Q-loop lid upon turnover, as shown here for *Ecbd*. This conformational change might be present only transiently, or require additional binding partners, prohibiting direct structural interrogation. Alternatively, we cannot exclude that short Q-loop *bd* oxidases have evolved a fundamentally distinct catalytic mechanism, and therefore inhibition with AurD, explaining why no quinone or AurD binding in these enzymes has been observed(*14, 30, 39, 40*).

The series of cryo-EM structures presented here enables us to propose a detailed mechanism for the entire turnover mechanism (Fig. 6). Even though our quinone bound structures represent the product rather than the substrate, we postulate that the minor difference in substrate (quinol) and product (quinone), do not result in major structural changes and therefore represent either state. We cannot exclude, however, that the equilibria between the different states change depending on the binding of either quinone or quinol. The catalytic cycle starts in the unbound state, in which the Q-loop remains flexible, and Tyr243^CydA^ is positioned for quinol capture (Fig. 6A). After quinol binding (Fig. 6B), the Q-loop undergoes a conformational transition that closes the lid and completes the active site by forming a hydrogen bond between Arg298^cydA^ and the quinone head (Fig. 6C). Proton-coupled electron transfer proceeds with electrons relayed through the heme chain to the oxygen reduction site at heme *d* (Fig. 6D), after which the Q-loop opens and the quinone is released (Fig. 6E). A second sequence of quinol oxidation occurs, during which O_2_ is fully reduced to H_2_O, completing the *Ecbd* turnover cycle.

**Figure 6.**
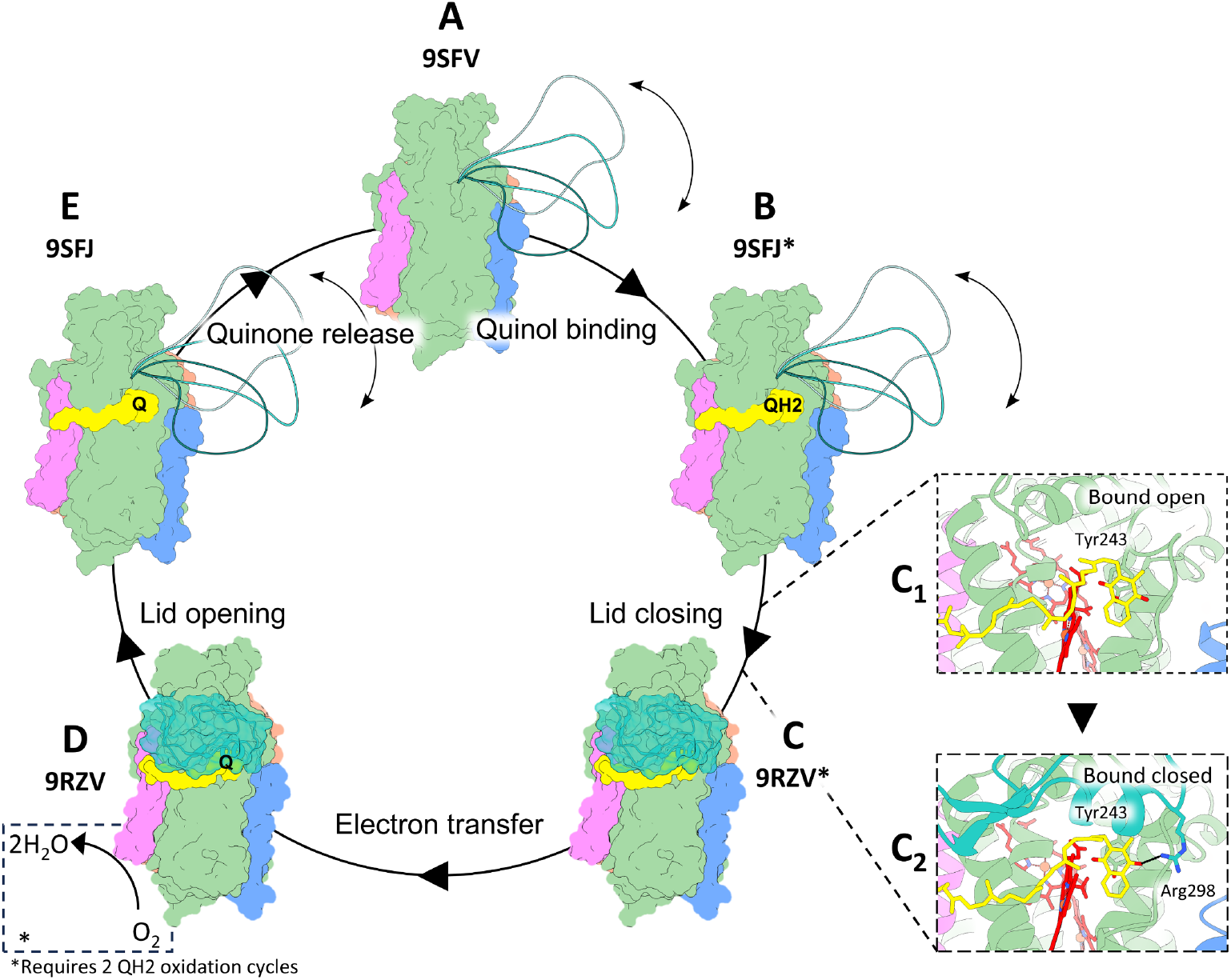
Schematic overview of the proposed turnover mechanism of *Ecbd*. (**A-D**) Overall schematic view of *Ecbd* turnover. States with an asterix are presumed to be structurally the same in the quinol and quinone bound states. (**A**) The unbound *Ecbd* dimer has a flexible Q-loop lid and sits ready for quinone binding. (**B**) The *Ecbd* dimer undergoes quinol binding. (**C**) From the quinone bound state (**C1**), the Q-loop lid closes, finishing the formation of the active site by coordinating Arg298 to the quinol head (**C2**). (**D**) electron transfer occurs via the heme chain onto molecular oxygen. (**E**) the Q-loop lid opens to release the quinone and start another catalytic cycle

Together, these insights provide a detailed molecular blueprint of *Ecbd* function and establish a foundation for rational drug design targeting *bd*-type oxidases. Given the essential role of cyt *bd* in the survival of several clinically relevant pathogens, these findings have broad implications for the development of next-generation antibiotics.

## Abbreviations

Cyt *bd*: Cytochrome *bd*
*Ecbd*: *E. coli* cytochrome *bd*-I
*Ecbd*-II: *E. coli* cytochrome *bd*-II
LMNG: Lauryl Maltose Neopentyl Glycol
MK: Menaquinone
NDH-2: NADH dehydrogenase type-2
cryo-EM: electron cryo-microscopy
CTF: contrast transfer function
NCC: normalized cross correlation

## Data availability

Cryo-EM maps are deposited at the Electron Microscopy Data Bank under accession codes: EMD-54414, EMD-54801, EMD-54812, EMD-54822, EMD-54823, EMD-54826, EMD-54866. Atomic models of *Ecbd* have been deposited to the Protein Data Bank under accession numbers: 9RZV, 9SE4, 9SEJ, 9SFF, 9SFH, 9SFJ and 9SFV. All other data are presented in the main text or supplementary information.

## Acknowledgements

We would like to thank Dr. Dirk Bald (VU Amsterdam, The Netherlands) for providing the MB43 and MB43ΔCydA cells for mutant expression, and Anneloes Cramer Blok for assisting with the SEC-MALS measurements. This work benefited from access to the Netherlands Center for Electron Nanoscopy (NeCEN) and the ALICE compute resources provided by Leiden University. Research reported in this publication was supported by Oncode Accelerator, a Dutch National Growth Fund project under grant number NGFOP2201, and by the Netherlands Electron Microscopy Infrastructure (NEMI), project number 184.034.014 of the National Roadmap for Large-Scale Research Infrastructure of the Dutch Research Council (NWO).

## Author contributions

L. J., T. T. v. d. V. conceptualization; S. B., K. K., T. T. v. d. V., and L. J. methodology; K. K. and T. T. v. d. V. formal analysis; F. P. and T. T. v. d. V. investigation; T. T. v. d. V. writing– original draft; S. B., K. K., and L. J. writing–review & editing; T. T. v. d. V. visualization; S. B. and L. J. supervision; L. J. project administration.

## Conflict of interest

The authors declare that they have no conflicts of interest.

## Experimental section

### Structure guided mutagenesis

Structure guided *Ecbd* mutagenesis was performed on the pET17b-CydABX-linkerstreptag(*34*) plasmid using Whole Plasmid Synthesis or Gibson assembly (See SI for primers). Mutants were confirmed using Sanger sequencing.

### *Ecbd* purification

The expression of *E. coli* cytochrome *bd*-I (*Ecbd*) was performed as stated previously(*15*). Briefly, MB43 cells(*41, 42*) transformed with pET17b-CydABX-linkerstreptag or MB43ΔCydA(*15*) transformed with the mutant plasmids were grown overnight in LB with 100 μg/mL ampicillin (250 RPM, 37°C). The WT culture was diluted to OD ∼0.1 and grown to OD∼ 0.4 before induction with 0.45 mM IPTG. At OD 2.0 the cells were harvested by centrifugation (6371 rcf, 20 min, 4°C) and resuspended in 50 mM MOPS, pH 7.4, 100 mM NaCl, cOmplete™ EDTA-free Protease Inhibitor (ROCHE), at 5 mL per 1 gram of wet cells. The *Ecbd* mutants were expressed overnight and otherwise followed the same procedure. The cells were lysed by a passing through a Stansted pressure cell homogeniser (270 MPa). Cell debris were pelleted by centrifugation (10.000 rcf, 20 min, 4°C). Crude membranes were isolated by ultra-centrifugation (200.000 rcf, 1h, 4°C) followed by resuspension in 50 mM MOPS, 100 mM NaCl, pH 7.4 (10 mg/mL total protein). Solubilization of the *Ecbd* was performed by incubation with 0.5% Lauryl maltose neopentyl glycol (LMNG) for 1 hour at 4°C with gentle mixing. Insoluble material was pelleted by ultra-centrifugation (200.000 rcf, 30 min, 4°C) followed by application of the soluble fraction to a StrepTrap HP column (Cytiva) at 2 mL/min. The column was washed with 50 mM sodium phosphate, 300 mM NaCl, 0.005% LMNG, pH 8.0 to remove unbound proteins. *Ecbd* elution was performed by addition of 50 mM sodium phosphate, 300 mM NaCl, 2.5 mM desthiobiotin, 0.005% LMNG, pH 8.0, after which purity was confirmed by SDS-page. If needed, *Ecbd* dimers and monomers were separated using size exclusion chromatography (SEC) on a Superdex increase 200 10/300 column (Cytiva) at 0.5 mL/min (50 mM sodium phosphate, 300 mM NaCl, 0.005% LMNG, pH 8.0). The defined fractions were pooled, concentrated, and stored at -80°C until further use.

### NDH2 purification

NDH2 was required to reduce the quinone pool in order to measure *Ecbd* activity. Expression and purification of NDH2 were performed as previously described(*22*). Briefly, C41 (DE3) cells, transformed with pET28-NDH-2_NtermHis, were grown overnight LB Kanamycin (250 RPM, 37°C). The culture was diluted 20-fold and grown to ∼OD 0.5 before induction with 0.25 mM IPTG. NDH2 expression was maintained for 4 hours at 37°C before harvesting by centrifugation (6371 rcf, 20 min, 4°C). The cells were resuspended in a 5 mL of 50 mM Tris-HCl, 5 mM MgCl_2_, pH 8.0 per 1 gram of cells, and disrupted by passing through a Stansted pressure cell homogeniser (270 MPa). Cell debris were pelleted by centrifugation (10.000 rcf, 20 min, 4°C) before harvesting of the crude membranes by ultra-centrifugation (200.000 rcf, 1h, 4°C). The membranes were resuspended in Tris-HCl, 150 mM NaCl, 20 mM Imidazole (10 mg/mL total protein). NDH2 was extracted by treatment with 1% DDM for 1 hour at 4°C with gentle mixing. The remaining insoluble material were removed by ultra-centrifugation (200.000 rcf, 30min, 4°C) before application of the soluble fraction on a HiTrap Nickel NTA column (Cytiva). The unbound proteins were washed from the column with 50 mM Tris-HCl pH 8.0, 150 mM NaCl, 20 mM Imidazole, 0.02% DDM. NDH2 was eluted by addition of 30% elution buffer (50 mM Tris-HCl, 150 mM NaCl, 500 mM Imidazole, 0.02% DDM). Final purification was achieved by gel filtration on a Superdex increase 200 10/300 column (Cytiva) at 0.5 mL/min (50 mM Tris-HCl, 500 mM NaCl, 5% glycerol, 0.02% DDM). Pure NDH2 fractions were pooled, concentrated, and stored at -80°C until further use.

### MSP1D1 expression and purification

MSP1D1 expression was performed as described before(*43*). Briefly, BL21(DE3) pLysS transformed with pET28a containing the MSP1D1 gene(*43*) were grown in terrific broth at 37°C, 200 RPM until OD ∼2.0. The cultures were cooled to 30°C before induction with 1 mM IPTG. The cells were induced for 5 hours, followed by cell harvesting (10.000 rcf, 20 min, 4°C). The cells were resuspended in 50 mM Tris-HCl (pH 8.0), 300 mM NaCl, 1% Triton X-100. The cells were lysed by a single pass through a Stansted pressure cell homogeniser (270 MPa). Debris and membrane fractions were pelleted by ultra-centrifugation (200.000 rcf, 1h, 4°C). The supernatant was applied to a HisTrap (Cytiva) column followed by extensive washing with buffer containing 1% (w/v) Triton X-100 and 50 mM Na-cholate. Finally, the MSP1D1 was eluted using 50 mM Tris, 300 mM NaCl, 300 mM Imidazole. The MSP1D1 peak fractions were concentrated and further purified on a HiLoad 16/100 75 pg column (Cytiva) using 50 mM Tris-HCl, pH7.5, 200 mM NaCl. The peak fractions were concentrated to 10 mg/mL and stored at - 80°C until further use.

### *Ecbd* reconstitution in nanodiscs

*Ecbd* nanodiscs were assembled as described previously with slight modifications(*13*). Briefly, the POPC:MK-9 mixture was dissolved in 20 mM HEPES (pH 7.4), 150 mM NaCl 100 mM Na-cholate by sonication. Following, MSP1D1, POPC, MK-9 and *Ecbd* were mixed at the molar ratio of 20:760:40:1 and incubated for 1h on ice. The detergent was removed by the stepwise addition of 5% w/v SM-2 biobeads every two hours, totaling 15% w/v before overnight incubation at 4°C while mixing. The biobeads were removed by filtration before application of the nanodisc mixture on a Superdex increase 200 10/300 column (Cytiva) at 0.5 mL/min (20 mM HEPES, 150 mM NaCl, pH 7.4). *Ecbd* nanodisc peak fractions were concentrated and used for further analysis and grid preparation.

### Nanodisc SEC-MALS analysis

The *Ecbd* nanodiscs were characterized using a SEC-MALS system comprised of a miniDAWN® TREOS®, Optilab differential refractometer (Wyatt technology) and 1260 Infinity II multiple wavelength absorbance detector (Agilent). The nanodisc composition was determined using the protein conjugate method in Astra software (Version 8) after defining the dn/dc_protein_, dn/dc_lipids_ and ε_417nm_, protein.

### *Ecbd* oxygen consumption

Oxygen consumption of *Ecbd* was measured on an oxygraph (Hansatech Ltd.) system at 20°C(*22*). The quinone (MK-1 or UQ-1) was added to the reaction chamber at the indicated concentration in 50 mM MOPS, 150 mM NaCl, (0.005 LMNG if needed) pH 7.0. The enzymatic quinone reduction with *C. thermarum* NDH-2 (30 nM) was initiated by the addition of 1 mM NADH, followed by the determination of the quinone autooxidation (background). Oxygen consumption was initiated by the addition of *Ecbd* (4 nM). The enzyme activity was measured by subtraction of the quinone autooxidation rate from the initial slope after *Ecbd* addition. In case of IC50 determination, the desired amount of AurD was added after Ecbd addition. The slope after AurD addition was divided by the slope before AurD addition to determine the inhibition rate.

### *Ecbd* reconstitution in proteoliposomes for electrochemistry

Lipids dissolved in chloroform were purchased from Avanti Polar Lipids. A lipid mixture of POPE:POPG:CA 60:30:10, enriched with the 1 mol% of ubiquinone-10 (UQ-10) (Sigma) or menaquinone-9 (MK-9) (Caymen chemical), was dried under a stream of nitrogen followed by overnight incubation under vacuum. The lipids were rehydrated to a final concentration of 10 mg/mL (20 mM MOPS, 30 mM Na_2_SO_4_, 100 mM KCl, pH 7.4) and extruded to 200 nm using an Avanti extruder. *Ecbd* reconstitution was performed as previously described(*26*). LMNG solubilized *Ecbd* was added to the liposome solution at 1 w/w% protein/lipids and mixed for 30 min by inversion at RT. Insoluble materials were removed by centrifugation in an Eppendorf tabletop centrifuge (14100 rcf, 5 min). The reconstituted *Ecbd* concentration was determined by re-dissolving a sample in 2% octyl-β-glucoside, followed by quantification of the Soret band (ε_417_ 230 mM^-1^ cm^-1^).

### Electrode preparation and electrochemistry in a tBLM system

Template-stripped gold (TSG) electrodes were produced in-house by evaporating a 150 nm thick 99.99% pure Au layer (Thessco) on a cleaned atomically smooth silicon wafer using a Plassys MEB600SL E-beam Evaporator operating at 10^-8^ mbar. The gold deposition was monitored using a piezoelectric quartz crystal at 6.0 MHz. Glass slides (1.2 cm^2^) were glued onto the gold layer with Epo-Tek 377 two-component glue and heated to 120°C for 2 hours for curing. Functionalization of the TSG electrodes with a self-assembled monolayer (SAM) was achieved by overnight incubation of freshly stripped electrodes in a mixture of 0.11 mM EO3-cholesteryl(*44*), 0.89 mM 6-mercaptahexanol (Sigma) dissolved in 2-propanol. Prior to measurement, the functionalized TSG electrodes were washed with 2-propanol and thoroughly dried under a stream of nitrogen. The TSG electrodes were placed into a custom polyether ether ketone (PEEK) sample holder with a Viton® O-ring seal (A = 0.29 cm^2^). The sample holder was placed into a custom electrochemical cell and submerged in the electrolyte solution (20 mM MOPS, 30 mM Na_2_SO_4_, pH 7.4). The electrochemical setup was completed by the addition of a Pt counter electrode, a saturated double junction Ag/AgCl (sat. KCl) reference electrode (Radiometer analytical) and placing it in a steel Faraday cage. All potentials are given versus standard hydrogen electrode (SHE) using 0.199 mV vs SHE for the Ag/AgCl reference electrode. The electrochemical measurement was performed using an Autolab (Eco Chemie) electrochemical analyzer equipped with a PGSTAT30 potentiostat and a FRA2 frequency analyzer. Impedance spectroscopy (EIS) was used to confirm that the SAM consisted of ∼50% EO3-cholesterol, as described previously(*45*). The electrolyte solution was purged with 95% Ar, 5% O_2_ to remove most oxygen. Cyclic voltammograms were taken at 10 mV/s by holding the potential at 0.4 V vs SHE for 5 seconds and scanning between 0.4 and -0.4 V vs SHE. Background cyclic voltammograms of the SAM were taken before the addition of the tethered bilayer lipid membrane (tBLM) system to the electrode. The tBLM was created by the addition of 10 mM CaCl_2_ and *Ecbd* proteoliposomes with 1 w/w% quinone (UQ-10 or MK-9) to a concentration of 0.5 mg/mL. The formation of the tBLM was followed by impedance spectroscopy and deemed finished once the signal stabilized (∼1 hour). The catalytic activity of *Ecbd* was measured using cyclic voltammetry.

### Cryo-EM Sample Preparation and Data Collection

Quantifoil UltrAUfoil R1.2/1.3 grids (mesh 300) were glow discharged with a PELCO easiGlow device at 15 mA for 90s on the foil side and 60 seconds on the other. *Ecbd* nanodiscs or in LMNG midelles (4 μL, 1.5 mg/ml), supplemented 100 with μM AurD if needed, were applied to the grid, followed by blotting at 4°C, 100% humidity, 20 blot force for 4 seconds using a Vitrobot IV device (Thermo Fisher) immediately before plunge freezing in liquid ethane. Blot time for *Ecbd* in LMNG was increased to 6 seconds.

Grids were either imaged on a Titan Krios G1 (ThermoFisher Scientific) operating at 300 kV, equipped with a Gatan K3 detector and BioQuantum energy filter with a slit width of 20 eV, or a Titan Glacios (ThermoFisher Scientific) operating at 200 kV with a Falcon 4i detector and selectris energy filter with a slit width of 20 eV. Movies were gathered in electron counting mode using aberration-free image-shift (AFIS) in EPU (ThermoFisher Scientific). A total dose of 100 e/Å2 with 100 frames, at 105,000× magnification with a calibrated pixel size of 0.836 Å (Krios), or a magnification of 130,000× with a calibrated pixel size of 0.880 Å (Glacios), at and a defocus range of 0.8 to 2.0 μm.

### Cryo-EM Data analysis

Data processing was performed in CryoSPARC. Image preprocessing was performed in CryoSPARC live(*46*) using Patch Motion Correction and Patch CTF estimation. The processed micrographs were exported to CryoSPARC for further processing. Initial particle picking was performed with blob picking, followed by template generation for both the monomer and dimer particles. Template particle picking was performed for the monomer and dimer particles followed by multiple rounds of 2D classification to remove junk particles. Three volume unsupervised ab initio model generation was performed to generate initial and bait volumes for further particle cleanup. Multiple rounds of heterogenous refinement was performed until the best class with clean particles became consistent. This clean particle stack was moved to Non-uniform refinement(*47*) for local CTF refinements and higher order aberration fitting. The movies were reprocessed in patch based motion correction to remove the last 50 frames to account for accumulated beam damage. The particles were subjected to reference based motion correction(*48*), followed by another round of Non uniform refinement before assessment of heterogeneity using 3D classification. Dimer particles in the apo dataset were split into the unbound and a bound-open class. The local resolution was improved using local refinement masking a single *Ecbd* protomer in the *Ecbd* dimer.

### Model building and analysis

Model building was started from pdb 6RKO for the *Ecbd* monomer and an alphafold(*49*) prediction for the dimer. Model building was performed in Coot software(*50*) (version 0.9.8.1) followed by real space refinement in Phenix(*51*) (version 1.21.1-5286). Models and their corresponding maps were assessed and analyzed in ChimeraX(*52*) (version 1.7.1). Interior cavities and tunnels were analyzed using MOLEonline(*28*) (5 probe, 1 interior threshold, 5 origin radius, 8 surface over radius, 1.1 bottleneck radius, 3 bottleneck tolerance).The pKa of the titratable residues along the proton channel were estimated using PROPKA. Ligand binding pockets were visualized using LigPlot+(*53*).

### Sequence alignment

For analysis of conserved residues, protein sequences were collected from the Interpro database for CydA (IPR002585), CydB (IPR003317), and CydX (IPR011724). This resulted in a database of 36K, 33K, and 5K sequences for CydA, CydB and CydX, respectively. Sequence alignment and analysis was performed using MAFFT.

## Notes

### Competing Interest Statement

The authors have declared no competing interest.

