## Supplementary information for "Visualizing the mechanism of quinol oxidation and inhibition of a *bd*-type oxidase using cryo-EM"

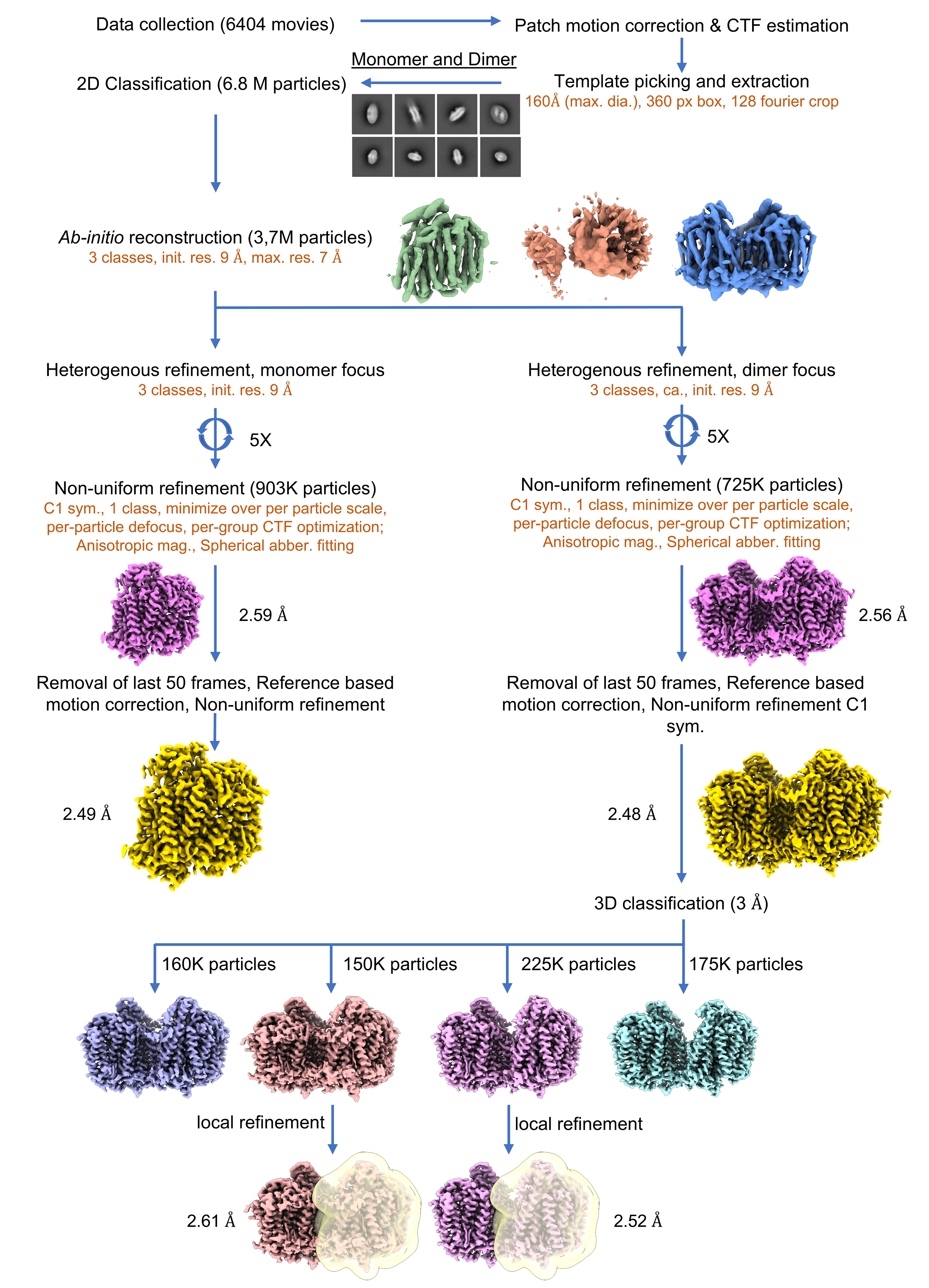

**SI Figure 1: Cryo EM processing of the Ecbd Apo structures**

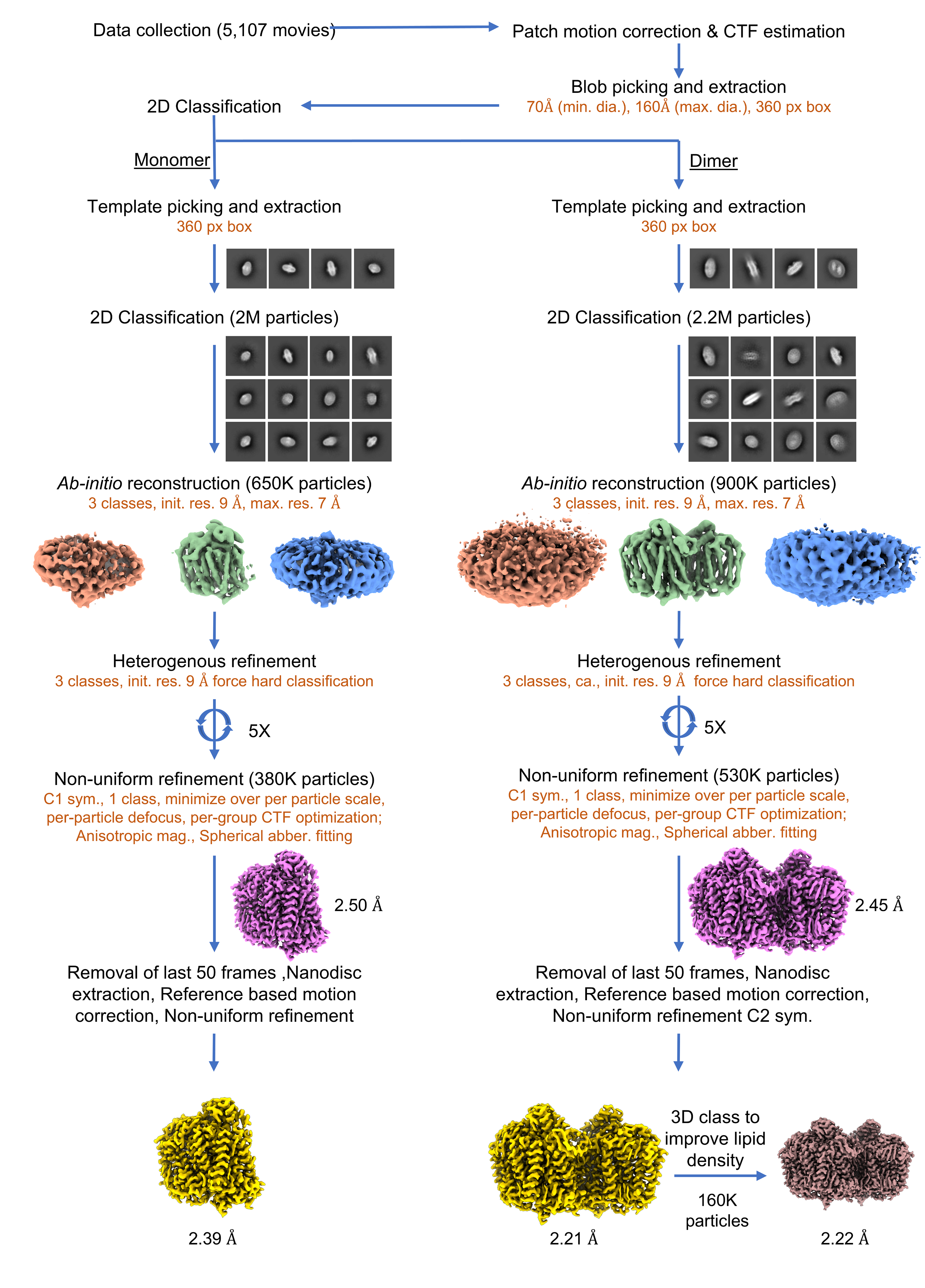

**SI Figure 2: Cryo-EM processing of the MK bound Ecbd structures**

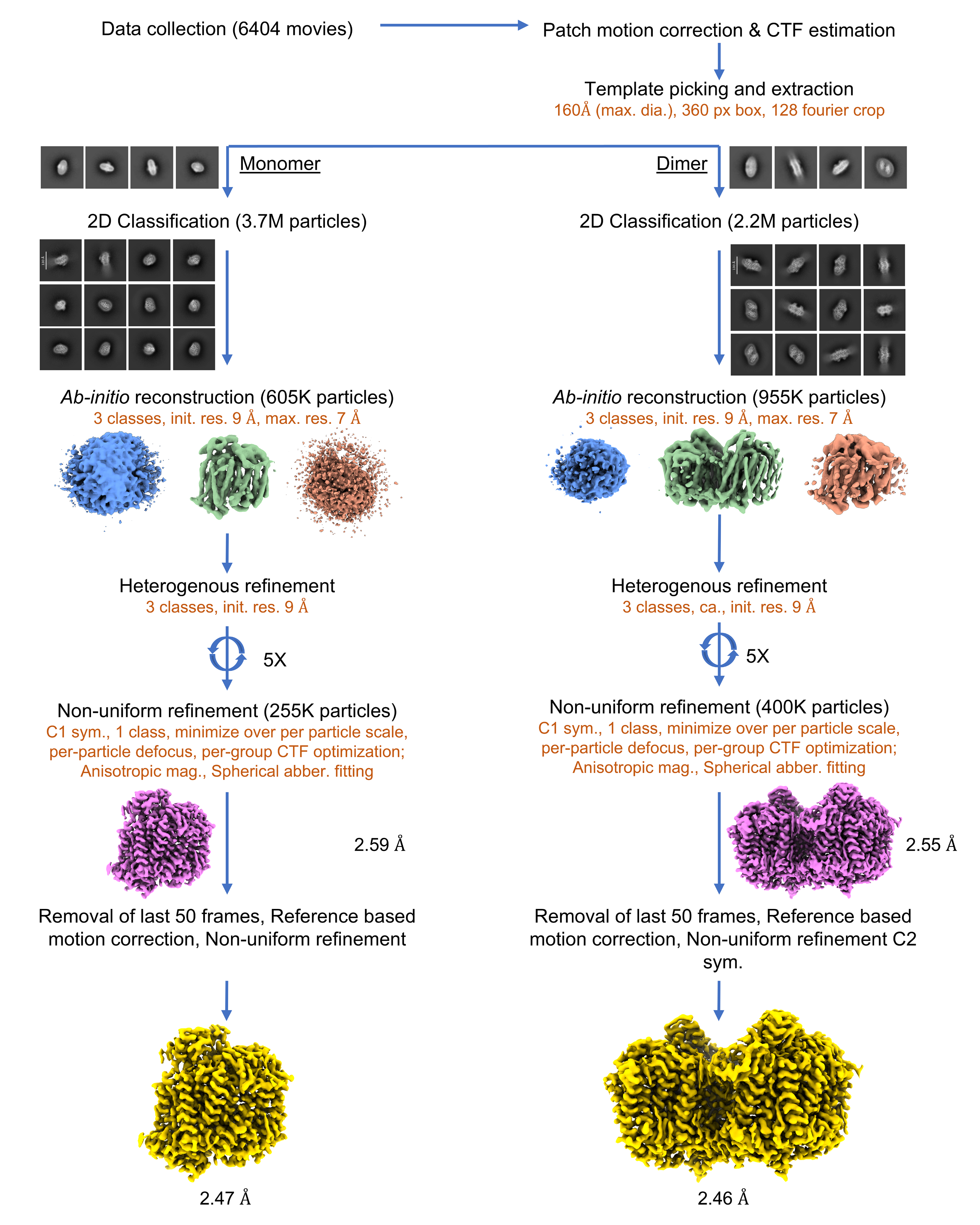

**SI Figure 3. Cryo-EM processing of the Aurachin D bound Ecbd structures.**

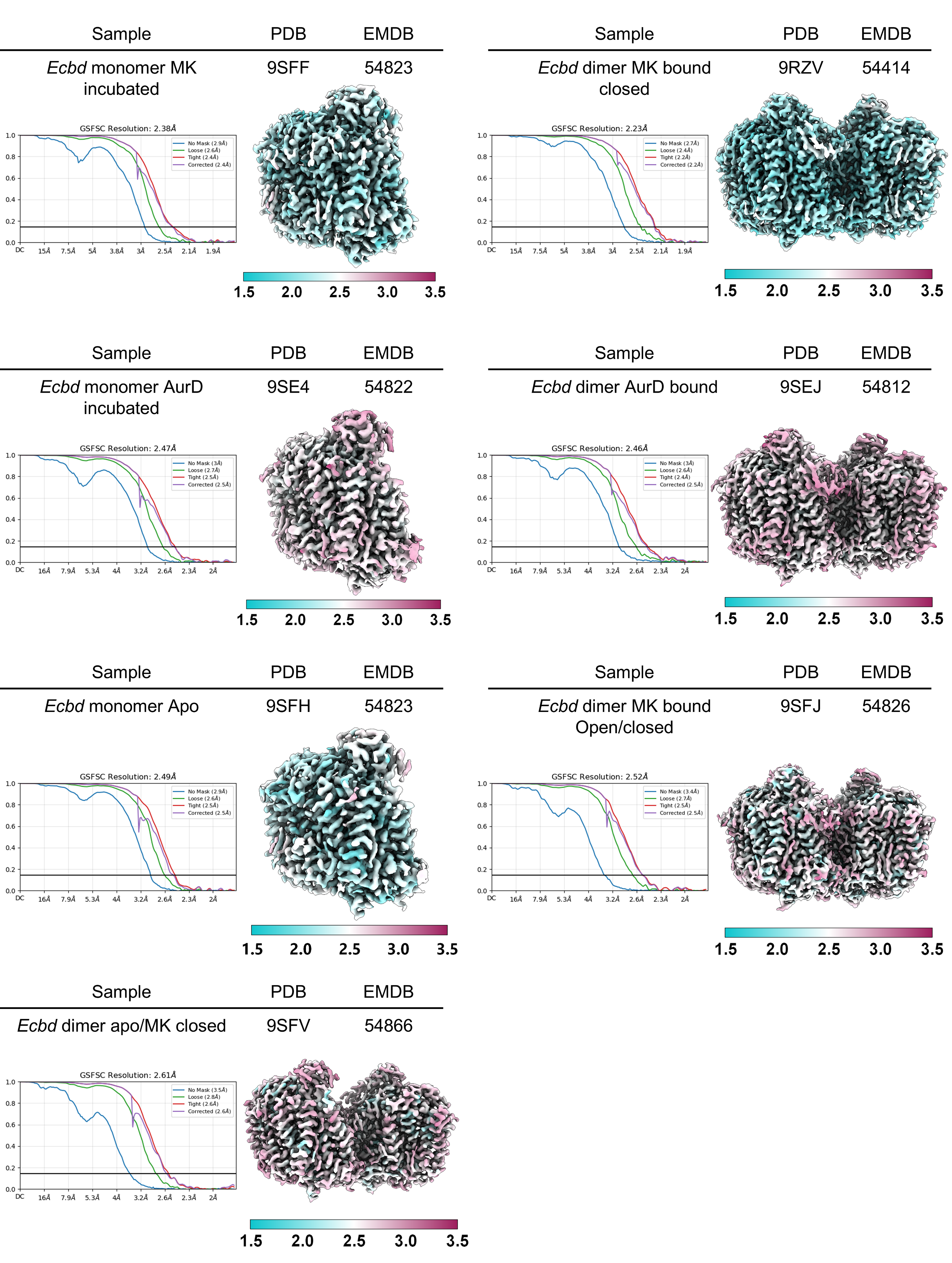

**SI Figure 4: GSFSC curves and local resolution estimates**

**Table 1. Cryo-EM data collection, refinement and validation statistics**

|  | ***Ecbd* monomer apo**  **(EMDB-54823)**  **(PDB 9SFH)** | ***Ecbd* monomer MK incubated**  **(EMDB-54822)**  **(PDB 9SFF)** | ***Ecbd* monomer AurD incubated**  **(EMDB-54801)**  **(PDB 9SE4)** | ***Ecbd* dimer MK bound Closed**  **(EMDB-54414)**  **(PDB 9RZV)** | ***Ecbd* dimer MK bound-open/closed**  **(EMDB-54826)**  **(PDB 9SFJ)** | ***Ecbd* dimer apo/MK closed**  **(EMDB-54866)**  **(PDB 9SFV)** | ***Ecbd* dimer AurD (EMDB-54812)**  **(PDB 9SEJ)** |
| --- | --- | --- | --- | --- | --- | --- | --- |
| **Data collection and processing** |  |  |  |  |  |  |  |
| Magnification | 130,000 | 105,000 | 130,000 | 105,000 | 130,000 | 130,000 | 130,000 |
| Voltage (kV) | 200 | 300 | 200 | 300 | 200 | 200 | 200 |
| Electron exposure (e–/Å^2^) | 100 | 100 | 100 | 100 | 100 | 100 | 100 |
| Defocus range (μm) | -0.8 - -2.0 | -1.0 - -2.2 | -0.8 - -2.0 | -1.0 - -2.2 | -0.8 - -2.0 | -0.8 - -2.0 | -0.8 - -2.0 |
| Pixel size (Å) | 0.880 | 0.836 | 0.880 | 0.836 | 0.880 | 0.880 | 0.880 |
| Symmetry imposed | C1 | C1 | C1 | C2 | C1 | C1 | C2 |
| Initial particle images (no.) | 6,873,607 | 2,979,789 | 3,721,318 | 3,289,126 | 6,873,607 | 6,873,607 | 2,009,440 |
| Final particle images (no.) | 764,760 | 380,310 | 399,575 | 530,094 | 224,083 | 152,474 | 255,127 |
| Map resolution (Å) | 2.49 | 2.39 | 2.47 | 2.23 | 2.52 | 2.61 | 2.47 |
| FSC threshold | 0.143 | 0.143 | 0.143 | 0.143 | 0.143 | 0.143 | 0.143 |
| Map resolution range (Å) | 2.46-2.6 | 2.39-2.51 | 2.47-2.56 | 2.21-2.28 | 2.52-2.58 | 2.5-2.56 | 2.45-2.51 |
| **Refinement** |  |  |  |  |  |  |  |
| Initial model used (PDB code) |  | 6RKO |  | Alphafold | | | |
| Model resolution (Å) | 2.6 | 2.4 | 2.5 | 2.2 | 2.4 | 2.5 | 2.3 |
| FSC threshold | 0.5 | 0.5 | 0.5 | 0.5 | 0.5 | 0.5 | 0.5 |
| Map sharpening *B* factor (Å^2^) | -60 | -60 | -60 | -40 | -60 | -60 | -60 |
| **Model composition** |  |  |  |  |  |  |  |
| Non-hydrogen atoms | 7492 | 7492 | 7492 | 16307 | 15464 | 15412 | 15975 |
| Protein residues | 892 | 892 | 892 | 1902 | 1902 | 1858 | 1902 |
| Ligands | 2 HEM, 1 A1JN4, 3 LPP, 1 UQ8, 1 OXY | 2 HEM, 1 A1JN4, 3 LPP, 1 MQ9, 2 POV, 1 OXY | 2 HEM, 1 A1JN4, 3 LPP, 1 UQ8, 1 OXY | 4 HEM, 2 A1JN4, 6 LPP, 4 MQ9, 4 PGT, 4 POV, 2 UQ8, 2 OXY | 4 HEM, 2 A1JN4, 6 LPP, 2 MQ8, 4 UQ8, 2 OXY | 4 HEM, 2 A1JN4, 6 LPP, 1 MQ8, 4 UQ8, 2 OXY | 4 HEM, 2 A1JN4, 4 LPP, 2 0NI, 2 PGT, 2 POV, 4 UQ8, 2 OXY |
| Water | 32 | 32 | 32 | 77 | 10 | 10 | 98 |
| *B* factors (Å^2^) |  |  |  |  |  |  |  |
| Protein | 68.40 | 79.60 | 63.40 | 53.92 | 97.41 | 124.62 | 78.28 |
| Ligand | 52.36 | 87.37 | 50.36 | 62.80 | 100.32 | 122.41 | 77.50 |
| **R.m.s. deviations** |  |  |  |  |  |  |  |
| Bond lengths (Å) | 0.003 | 0.003 | 0.003 | 0.004 | 0.004 | 0.004 | 0.004 |
| Bond angles (°) | 0.805 | 0.703 | 0.806 | 0.749 | 1.460 | 1.140 | 1.47 |
| **Validation** |  |  |  |  |  |  |  |
| MolProbity score | 1.60 | 1.39 | 1.58 | 1.31 | 1.40 | 1.32 | 1.37 |
| Clashscore | 7.79 | 5.68 | 7.51 | 2.01 | 5.05 | 2.29 | 4.28 |
| Poor rotamers (%) | 1.5 | 0.55 | 1.64 | 0.26 | 1.32 | 1.12 | 0.58 |
| **Ramachandran plot** |  |  |  |  |  |  |  |
| Favored (%) | 97.85 | 97.84 | 97.96 | 99.26 | 97.83 | 97.61 | 97.93 |
| Allowed (%) | 2.04 | 2.16 | 2.04 | 0.96 | 2.17 | 2.39 | 1.96 |
| Disallowed (%) | 0.11 | 0 | 0 | 0.05 | 0 | 0 | 0.11 |

**Table 2. Mutagenesis primers**

| **Primer** | **Sequence** |
| --- | --- |
| *Ecbd.A* S233A Fw | atggctgctgttctggctgttattgttctgggt |
| *Ecbd.A* S233A Rv | acccagaacaataacagccagaacagcagccat |
| *Ecbd.A* D239A Fw | tctgttattgttctgggtgccgaatccggctacgaaatgggc |
| *Ecbd.A* D239A Rv | gcccatttcgtagccggattcggcacccagaacaataacaga |
| *Ecbd.A* D239G Fw | tctgttattgttctgggtggcgaatccggctacgaaatgggc |
| *Ecbd.A* D239G Rv | gcccatttcgtagccggattcgccacccagaacaataacaga |
| *Ecbd.A* D239P Fw | attgttctgggtccagaatccggctac |
| *Ecbd.A* D239P Rv | gtagccggattctggacccagaacaat |
| *Ecbd.A* Y243A Fw | ctgggtgacgaatccggcgccgaaatgggcgacgtgcag |
| *Ecbd.A* Y243A Rv | ctgcacgtcgcccatttcggcgccggattcgtcacccag |
| *Ecbd.A* R298A Fw | ctgggcatcattgcaacggcatccgtggataccccggtt |
| *Ecbd.A* R298A Rv | aaccggggtatccacggatgccgttgcaatgatgcccag |
| *Ecbd.X* T10A Fw | gcatggattctgggagctcttcttgcctgttcg |
| *Ecbd.X* T10A Rv | Cgaacaggcaagaagagctcccagaatccatgc |
| *Ecbd* Gibson Fwd | gcccgaaaggaagctgagtt |
| *Ecbd* Gibson Rev | aactcagcttcctttcgggc |

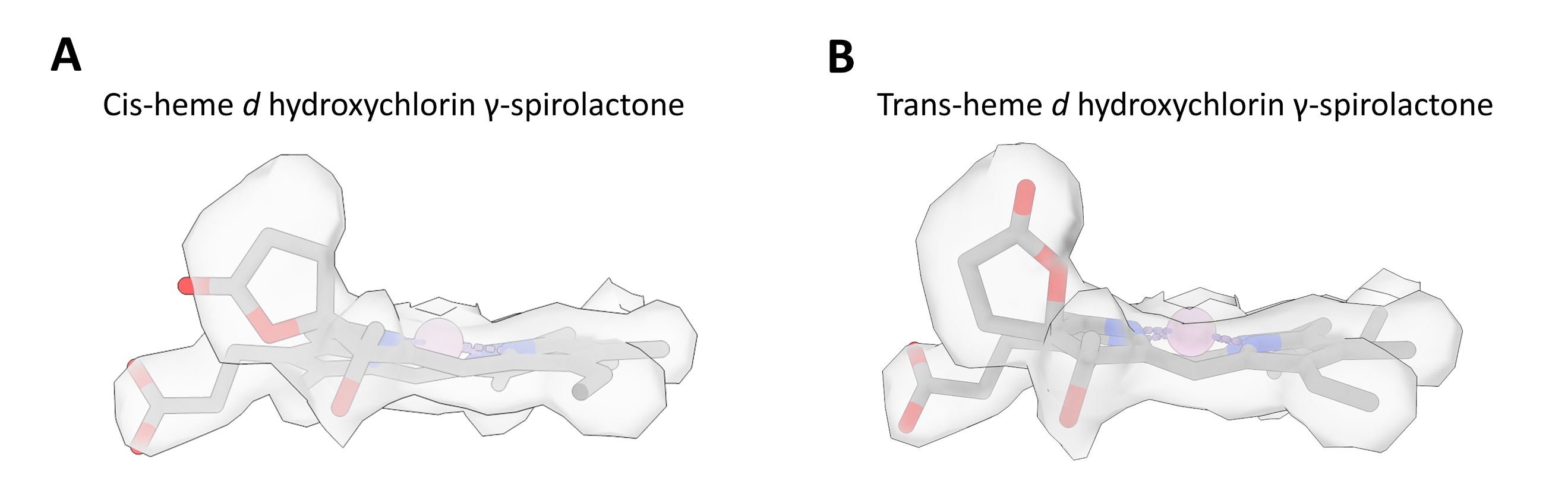

**SI Figure 5. Fitting of the heme density with** (**A**) cis heme d hydroxychlorin γ-spirolactone or (**B**) trans heme d hydroxychlorin γ-spirolactone

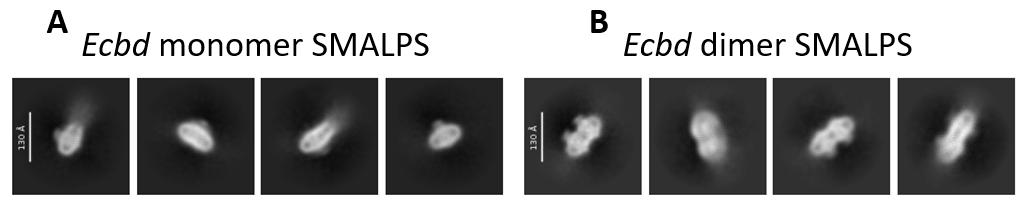

**SI Figure 6. 2D classes of Ecbd after SMALP isolation** (**A**) 2D classes of the Ecbd monomer (**B**) 2D classes of the Ecbd dimer showing of both oligomeric states in vivo.

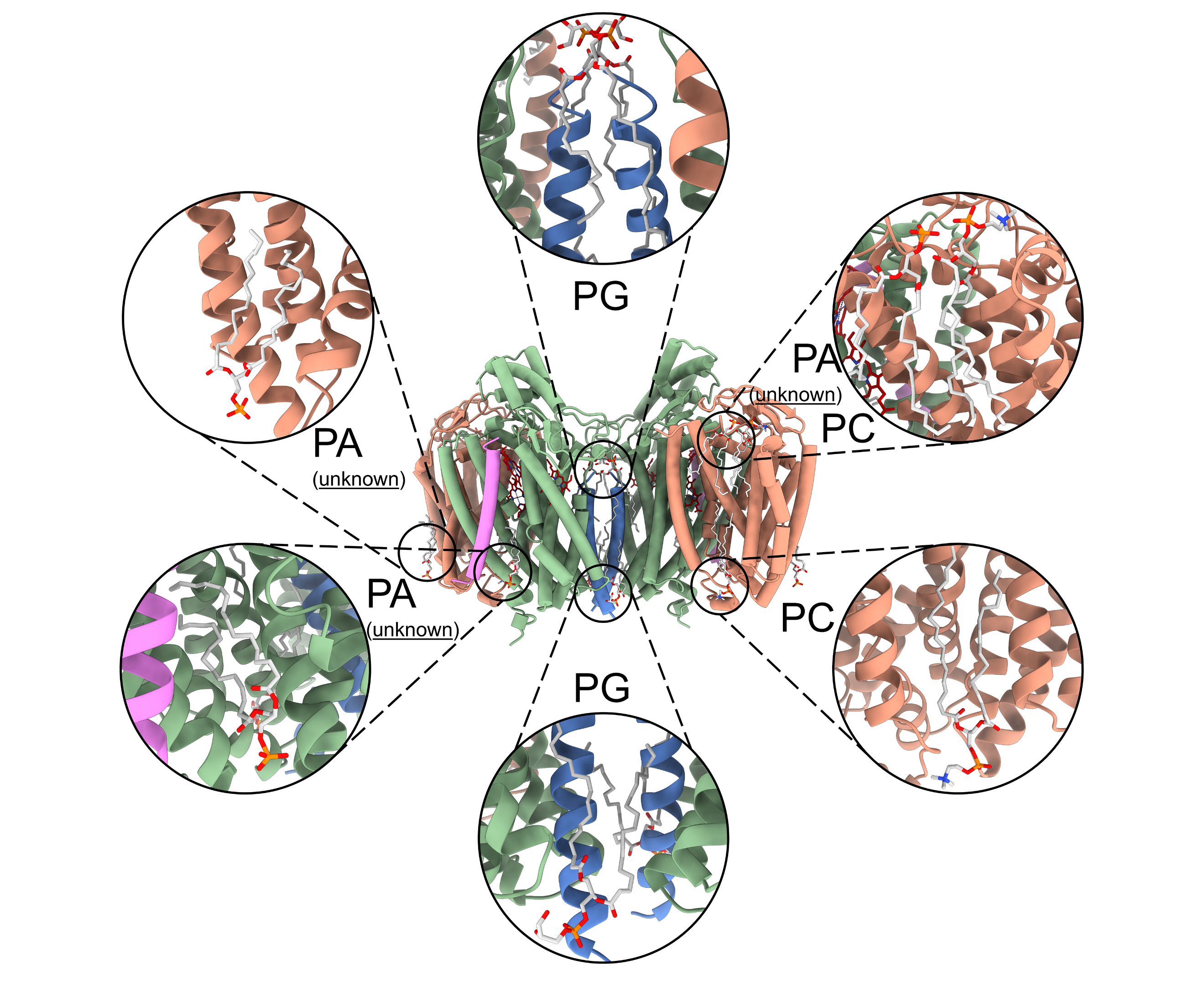

**SI Figure 7. Lipids bound peripherally and to the interface of the Ecbd dimer.** Lipids with unclear headgroup densities are modelled as phosphatidic acid (PA).

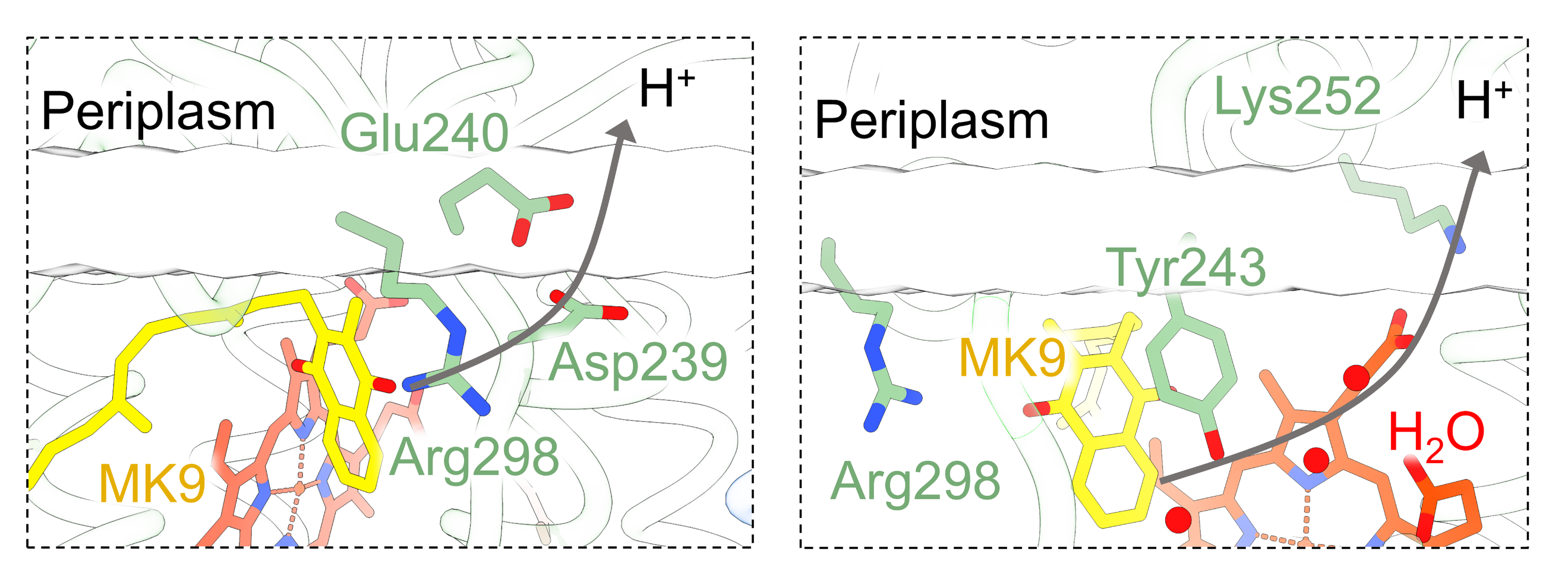

**SI Figure 8. Putative proton transfer routes from the quinol oxidation site towards the periplasm.** One route transfers past the Asp239 Glu240 pair near the quinol oxidation site. The other route transfers via structures water molecules towards Lys252 at the membrane interface.

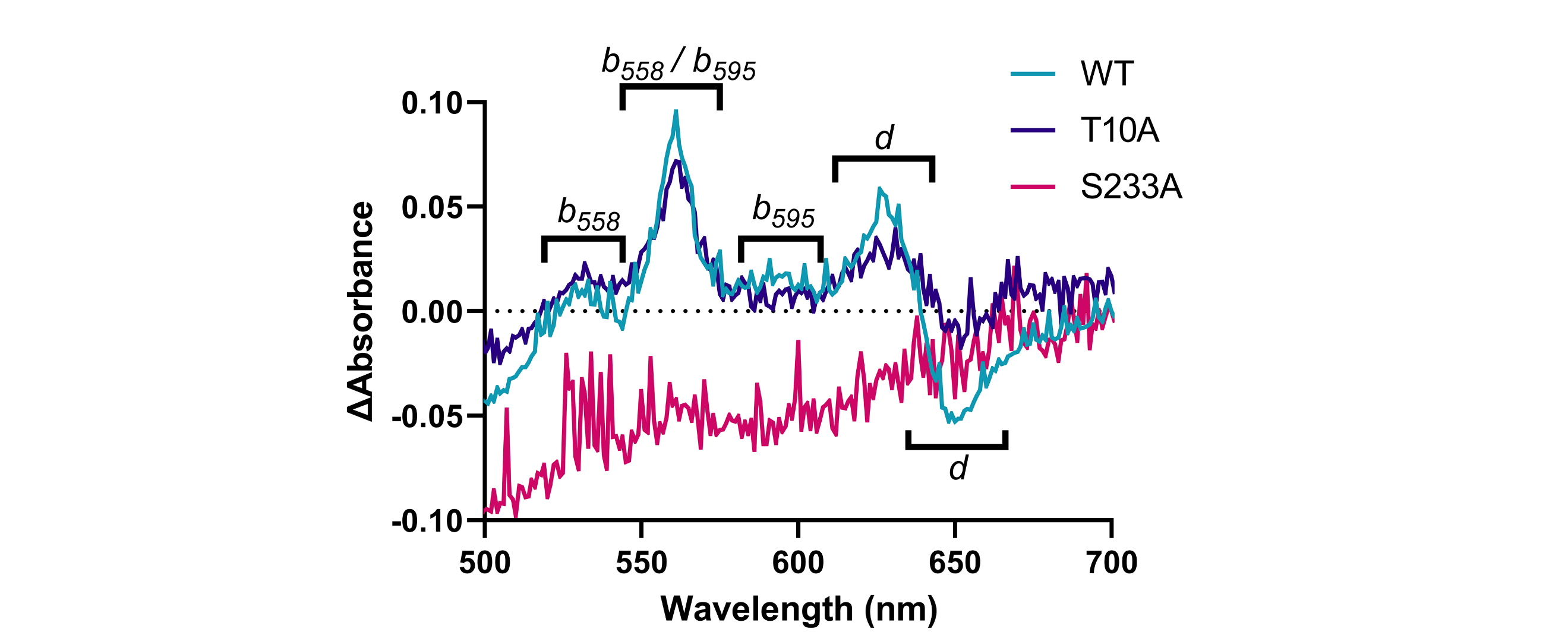

**SI Figure 9. Reduced minus oxidized spectra of the Ecbd WT and the T10A, S233A mutants**

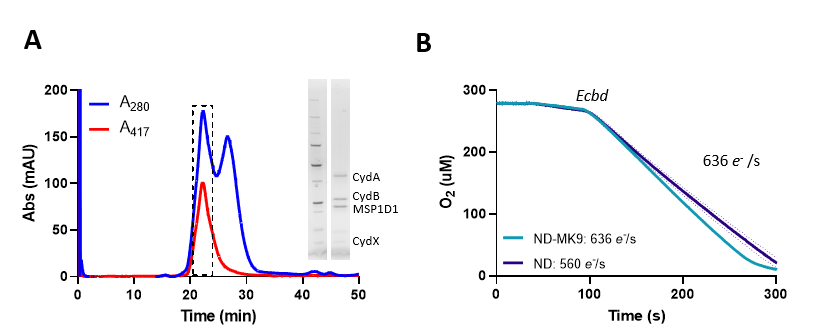

**SI Figure 10. Ecbd reconstitution in nanodiscs.** (**A**) Size exclusion of Ecbd nanodiscs. (**B**) Activity of Ecbd nanodiscs with either MK or UQ.

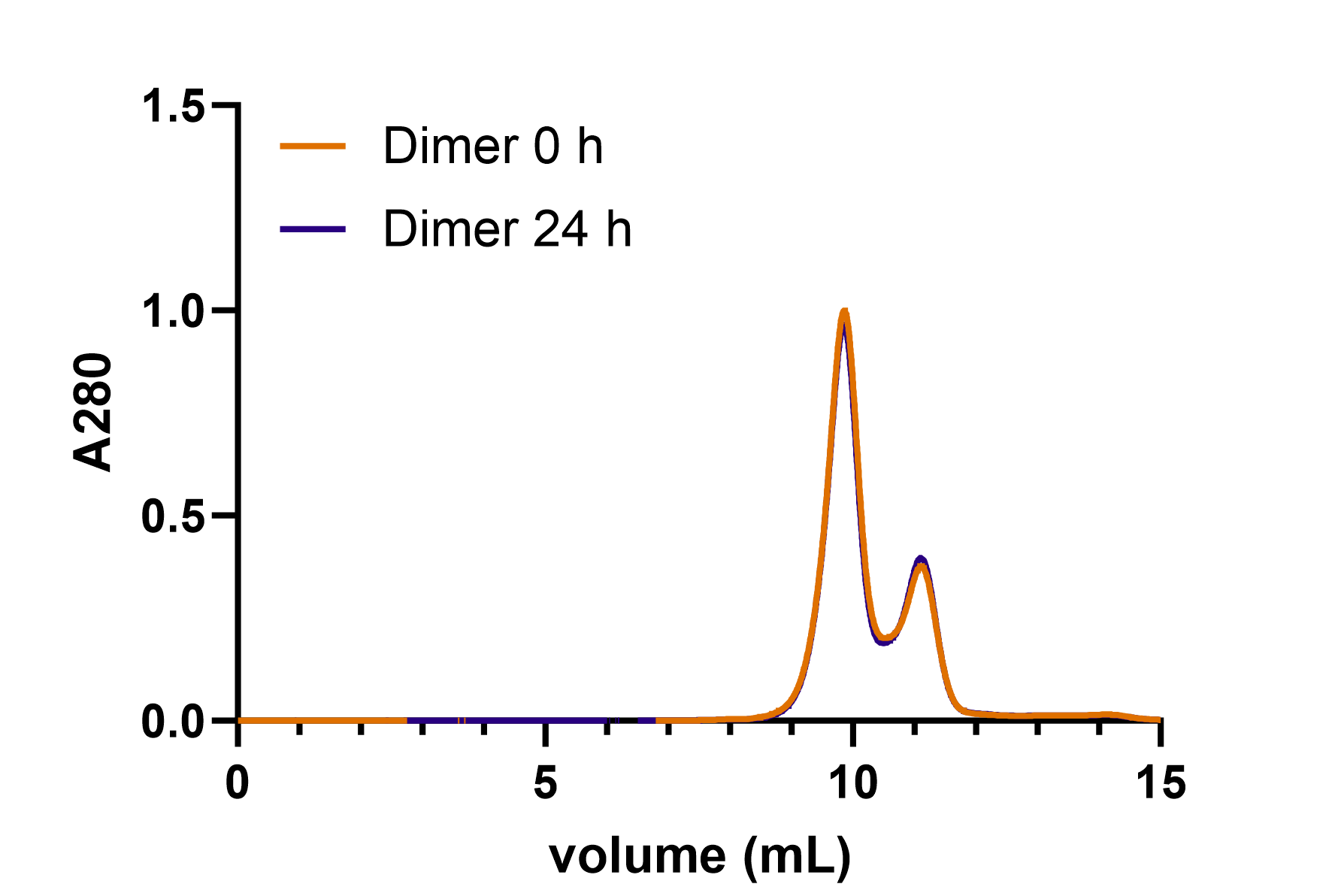

**SI Figure 11. SEC-MALS trace of the Ecbd dimer in detergent over 24 hours.**

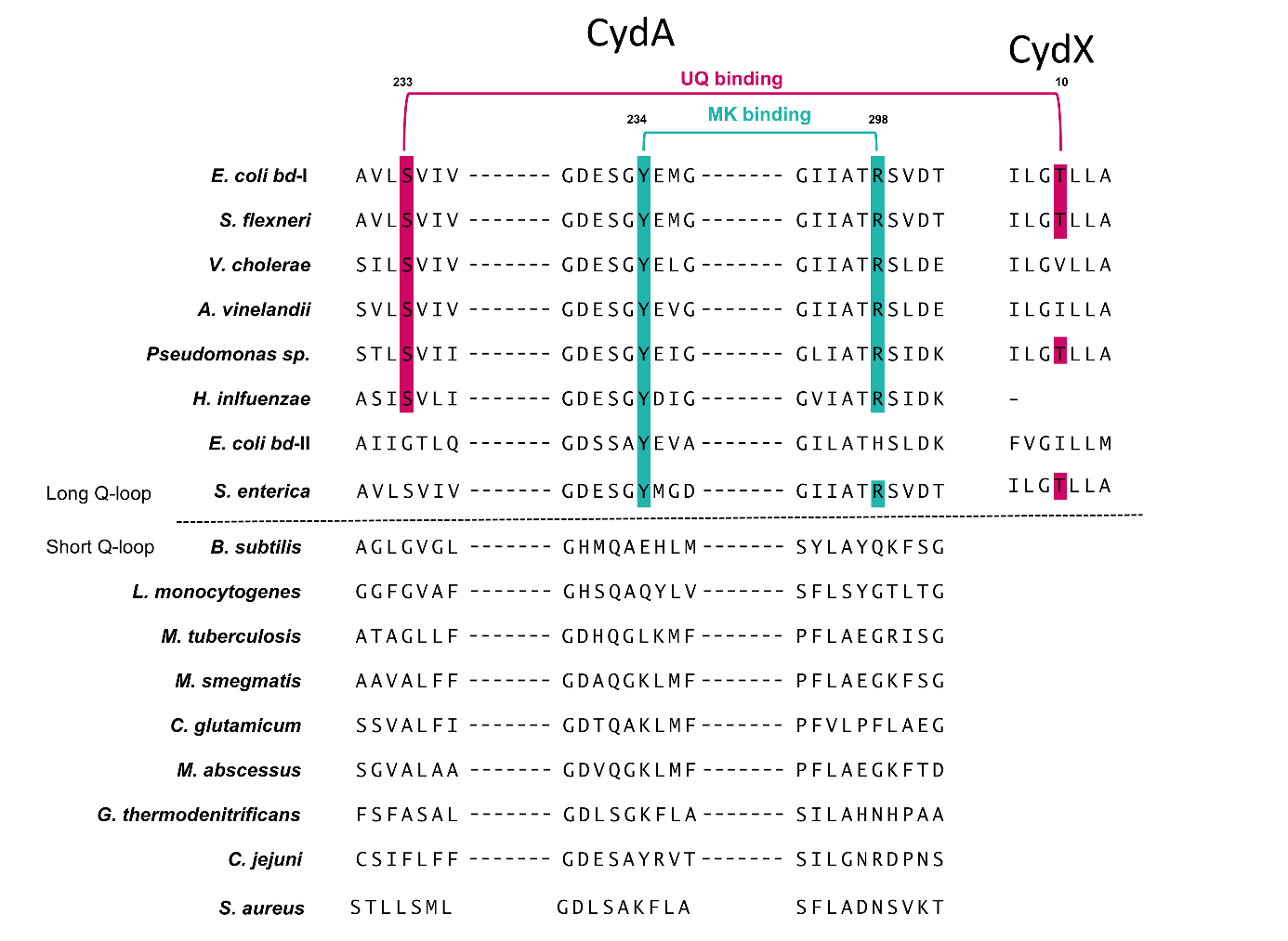

**SI Figure 12. Extended sequence alignment of CydA from different species across the long and short Q-loop families.**

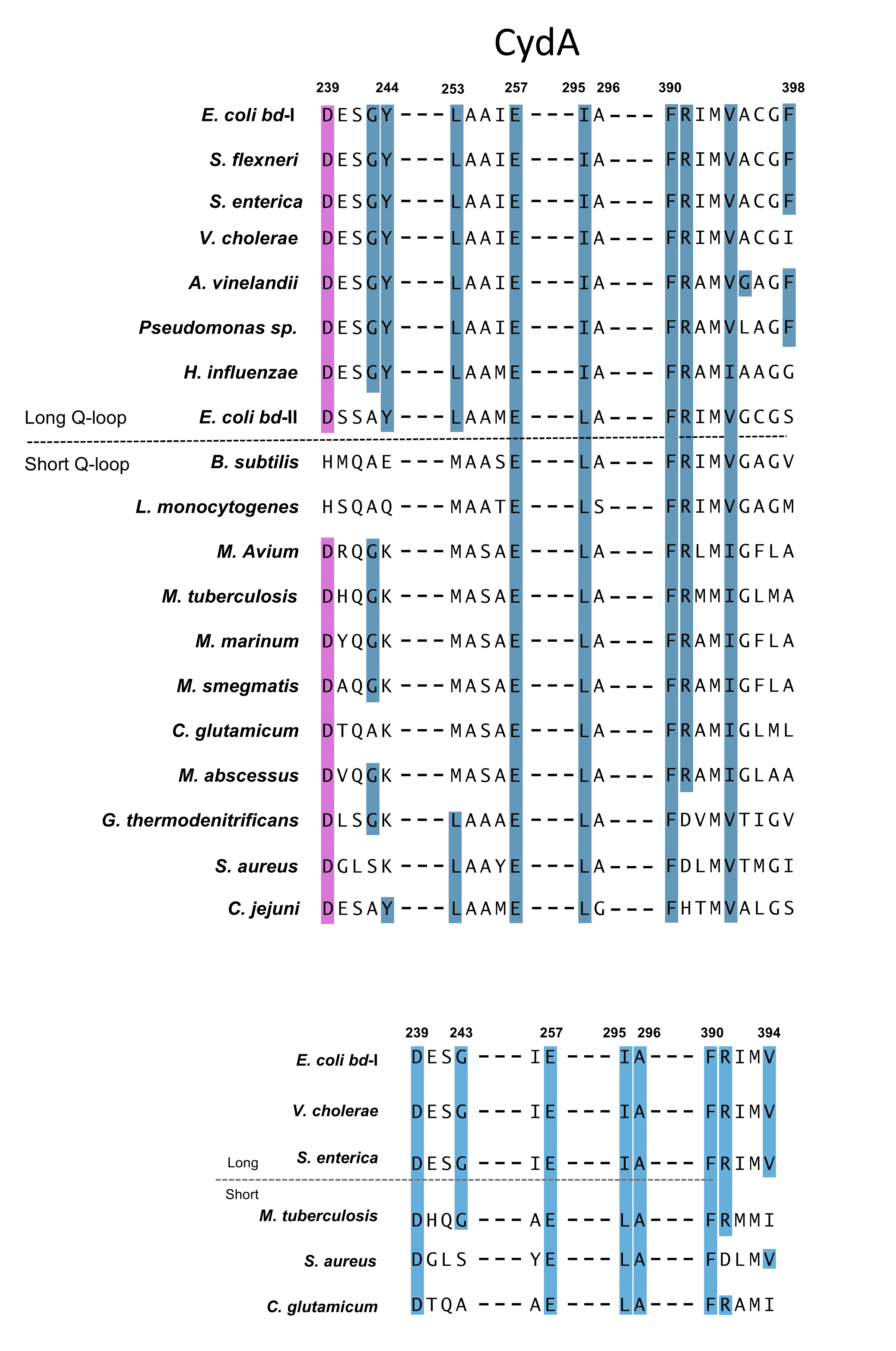

**SI Figure 13: Extended sequence alignment of the AurD binding pocket across bd oxidases**

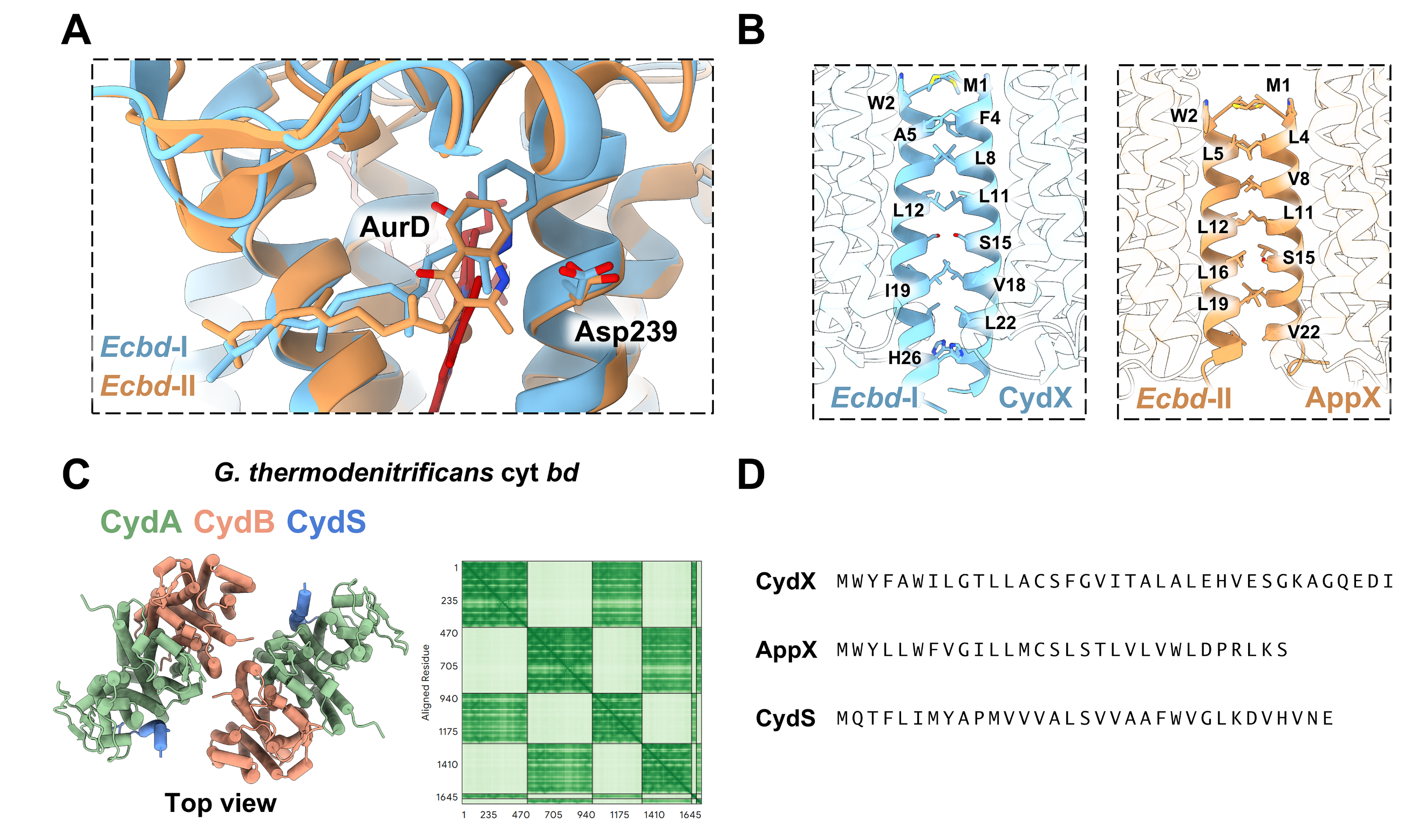

**SI Figure 14. Comparison of AurD binding and the dimer interface** (**A**) binding of AurD to Ecbd-I and Ecbd-II (7OSE) (**B**) dimer interface of Ecbd-I CydX and Ecbd-II AppX (**C**) Predicted G. thermodenitrificans cyt bd dimer using alphafold 3. (**D**) sequence comparison of E. coli CydX, AppX and G. thermodenitrificans CydS.

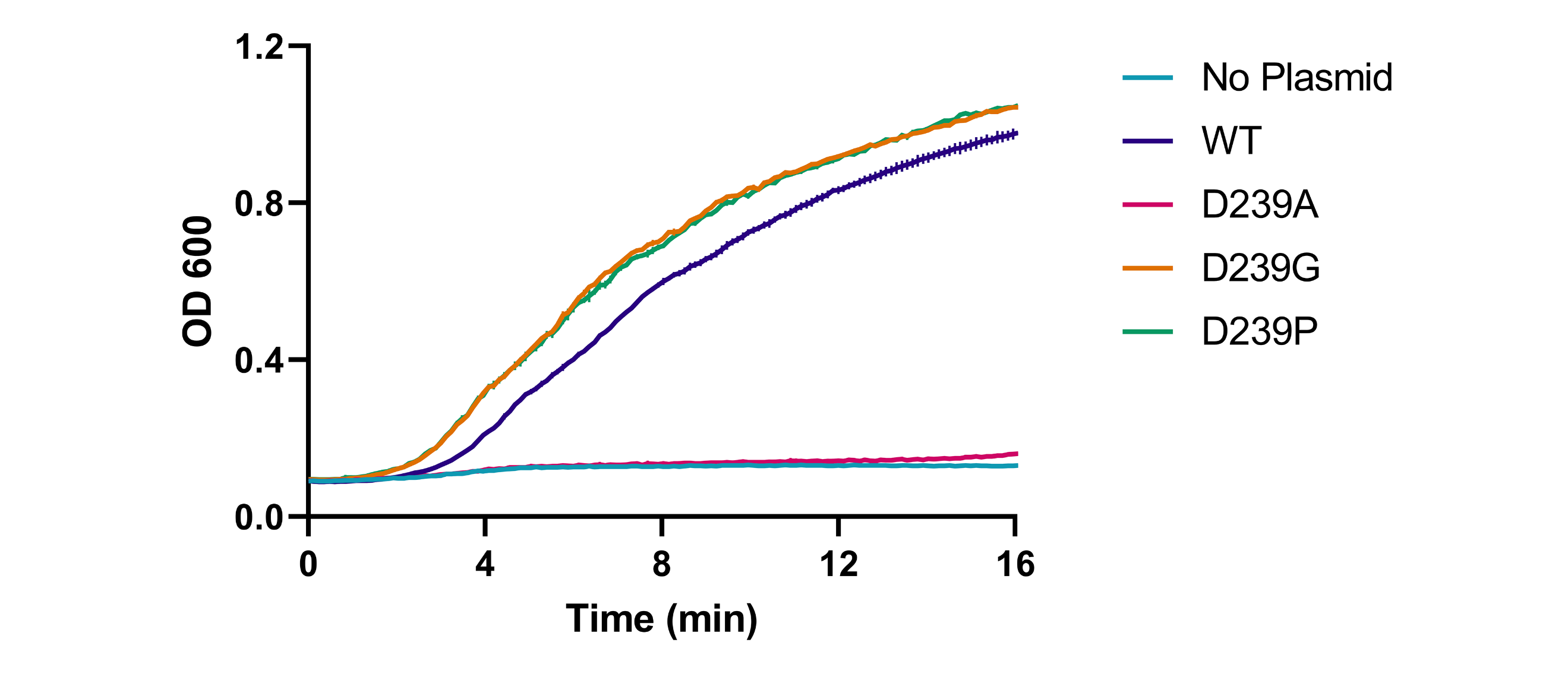

**SI Figure 15. Growth curves of the MB43 ΔcydA knockout strain supplemented with Ecbd WT or its D239 mutants**

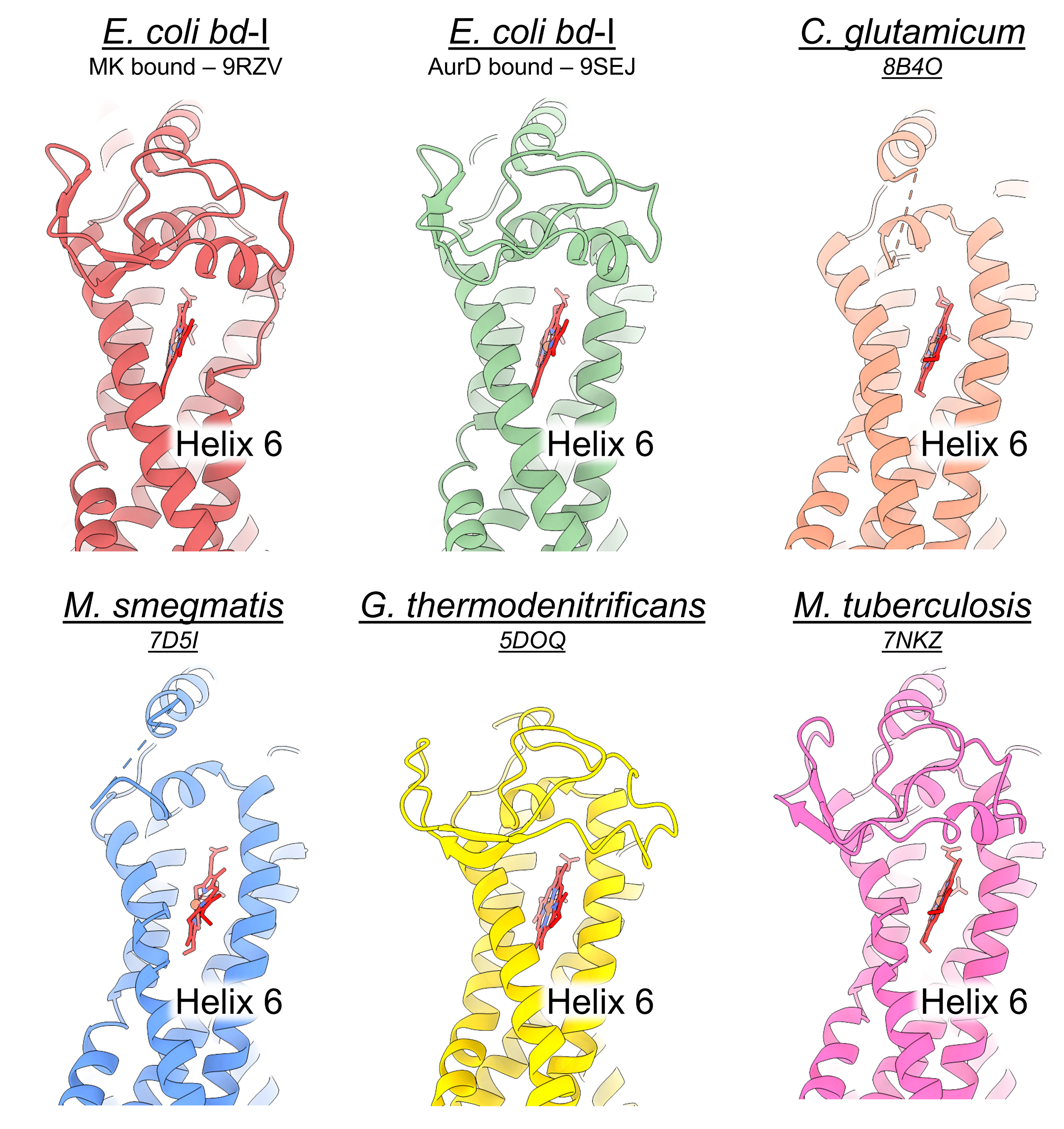

**SI Figure 16. Overview of the helical states of cyt bd oxidases.** The short Q-loop bd oxidases represent the helical fold of helix 6, as in the AurD inhibited state of E. coli cytochrome bd-I

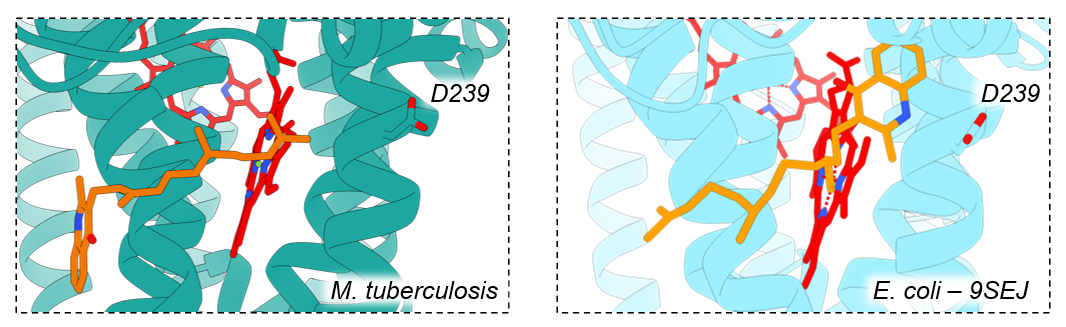

**SI Figure 17. Comparison of the predicted binding pose of Aurachin D in M. tuberculosis cytochrome bd using AutoDock Vina**(19, 20) **via Swissdock**(21, 22)**, with the cryo-EM structure of Ecbd bound to Aurachin D (9SEJ).**
